# Genome-wide identification and prediction of SARS-CoV-2 mutations show an abundance of variants: Integrated study of bioinformatics and deep neural learning

**DOI:** 10.1101/2021.05.23.445341

**Authors:** Md. Shahadat Hossain, A. Q. M. Sala Uddin Pathan, Md. Nur Islam, Mahafujul Islam Quadery Tonmoy, Mahmudul Islam Rakib, Md. Adnan Munim, Otun Saha, Atqiya Fariha, Hasan Al Reza, Maitreyee Roy, Newaz Mohammed Bahadur, Md. Mizanur Rahaman

## Abstract

Genomic data analysis is a fundamental system for monitoring pathogen evolution and the outbreak of infectious diseases. Based on bioinformatics and deep learning, this study was designed to identify the genomic variability of SARS-CoV-2 worldwide and predict the impending mutation rate. Analysis of 259044 SARS-CoV-2 isolates identify 3334545 mutations (14.01 mutations per isolate), suggesting a high mutation rate. Strains from India showed the highest no. of mutations (48) followed by Scotland, USA, Netherlands, Norway, and France having up to 36 mutations. Besides the most prominently occurring mutations (D416G, F106F, P314L, and UTR:C241T), we identify L93L, A222V, A199A, V30L, and A220V mutations which are in the top 10 most frequent mutations. Multi-nucleotide mutations GGG>AAC, CC>TT, TG>CA, and AT>TA have come up in our analysis which are in the top 20 mutational cohort. Future mutation rate analysis predicts a 17%, 7%, and 3% increment of C>T, A>G, and A>T, respectively in the future. Conversely, 7%, 7%, and 6% decrement is estimated for T>C, G>A, and G>T mutations, respectively. T>G\A, C>G\A, and A>T\C are not anticipated in the future. Since SARS-CoV-2 is evolving continuously, our findings will facilitate the tracking of mutations and help to map the progression of the COVID-19 intensity worldwide.

## 1. Introduction

The novel coronavirus (SARS-Cov-2) is the culprit for the ongoing pandemic of COVID-19 disease which is a respiratory like disease showing pneumonia-like symptoms and was first recorded in December 2019 in Wuhan, China ^1^. The single-stranded RNA genome of this infectious agent contains several ORF encoding structural as well as non-structural proteins. Spike glycoprotein (S), Envelop protein (E), Membrane protein (M), Nucleocapsid protein are the structural proteins whereas 3-chymotrypsin-like protease, papain-like protease and RNA-dependent RNA polymerase are the non-structural proteins ^2^.

Conscious of becoming an international health emergency, the World Health Organization (WHO) proclaimed COVID-19 as a pandemic on 11^th^ March 2020. From then the virus has now spread to over 223 nations and territories (https://www.who.int/emergencies/diseases/novel-coronavirus-2019/situation-reports). A disparity was found in cases of mortality in various countries, perhaps because of a different demographic makeup and the form of strategies adopted in different countries to restrict the transmission of the virus ^3^. According to the WHO 166,840,355 confirmed cases of Corona Virus Disease-19 (COVID-19) have been reported with 3,464,312 death worldwide by May 22, 2021 (https://www.worldometers.info/coronavirus/). The varying infection outcome, mechanism of transmission and time of incubation collectively have reinforced the capability of the virus to spread effectively across the world. This issue of the high virus transmissibility from one human to another is being constantly addressed by the researchers ^4^.

SARS-CoV-2 shows a higher mutation and evaluation rate than that of its hosts, making it able to adapt against the immune system within the host body. Mutation can be occurred due to certain errors in SARS-CoV-2 genome during the replication process, specifically the time of copying RNA to a new cell ^5^. Single nucleotide polymorphism (SNP), Insertion, and Deletion are the main type of mutations. Single nucleotide polymorphism (SNP) can be divided into silent/nonsense (synonymous SNP), missense (non-synonymous SNP), and frameshift subclasses ^6^. In viral evolution mutation rate is one of the critical parameters and the micro-level alteration in the mutation rate can change the characteristics of the virus and virulence dramatically in the host immune systems ^7^. A precise assessment of the mutation rate therefore becomes significant to assume the risk of emerging infectious diseases like SARS-CoV-2 ^8^.

SARS-Cov-2 has a strong nucleotide identity (96%) with bat coronavirus and for little dissimilarity. The virus may have gone through a process of adaptation and this provides opportunity to the virus to jump species boundaries and cause disease in humans ^2,9^. There was little evidence of local or regional adaptation throughout the initial phylogenomic study of three main clades (S, V, and G) of Global Initiative on Sharing All Influenza Data (GISAID) separated from outbreaks of multiple geographic locations (China, USA, and Europe) within SARS-CoV-2, proposing instead that viral evolution is driven solely by genetic drift and founding events. Even so, several reports anticipate possible nucleotide, amino acid (aa) adaptation, and structural heterogeneity in viral proteins, notably in spike protein ^10-12^. There are numerous deletions observed in the SARS-CoV-2 variants from different geographic locations. In a Singapore case cluster, a SARS-CoV-2 variant with a 382-nucleotide deletion was detected in January to February 2020 and also detected in a traveler who returned from Wuhan, China, to Taiwan in February 2020 ^13,14^, which is responsible for decollate the ORF7b and preventing the transcription of ORF8 by eliminating the transcription-regulatory sequence of ORF8 and this variant was not detected after the march, 2020 though it was successfully transmitted. ORF8 deletion in multiple SARS-CoV-2 isolates identified in the cases of Bangladesh (345 nucleotides), Australia (138 nucleotides), and Spain (62 nucleotides) ^14^. Unique or multiple SARS-CoV-2 clades or mutation(s) and clinical symptoms of COVID-19 observed between countries thus it is important to integrate sequence determination of related infectious genome with patient clinical knowledge (sequence polymorphisms and symptoms) to have the best understanding of SARS-CoV-2 pathogenicity ^1,9,15-17^. In addition, genomic sequence and mutation analysis are critical for the inventing of proper drugs and vaccines against this RNA virus ^18^. Indeed, reliable data on the rate of viral mutation can play a crucial role in evaluating potential vaccination strategies.

From this sense, we aimed to perform a large scale study over the SARS-CoV-2 genome to identify the base substitution mutations along with the rate of mutation using the available dataset in the GISAID. From GISAID, we have analyzed the complete genome sequence of 259044 viral samples isolated from different countries for a certain period of December 2019 to December 2020. We concentrate particularly on mutations which have freely evolved on different dates because these are the possible opportunities for further adaptation of SARS-CoV2 to its human host. Afterward, based on the findings from the mutational analysis we intended to predict the mutation rate of the virus for future times through the deep learning approach lied on artificial recurrent neural network (RNN) called Long Short Term Memory (LSTM). It is anticipated that the present study will contribute to understand the evolving nature of SARS-CoV-2 in the human body and will help to establish strategies to tackle the epidemiological and evolutionary levels.

## 2. Results

### 2.1 Overall mutational profile of SARS-CoV-2 genome

Worldwide a total number of 259044 complete genome sequence of SARS-CoV-2 were taken to investigate the overall genetic variations over the NC_045512.2 Wuhan reference genome and this analysis identifies a total of 3334545 mutations. Most of the samples possessed more than one mutation where 17221samples were found to contain 18 mutations followed by the 17, 16 and 19 mutations for 17084, 14983, and 14801 samples, respectively. The highest number of 48 mutations were observed for 1 sample (**Figure 1**). Average mutations per sample was calculated as 14.01. Among the 259044 complete genome sequence of SARS-CoV-2, the top 20 most mutated samples were identified, and we found that a sample from India had the maximum number of mutations (48). Samples from Scotland, USA, Netherlands, Norway, and France were found to have up to 36 mutations (**Figure 2**). After the onset of SARS-CoV-2 at Wuhan in China, 4 mutations were found in December 2019 (**Figure S1**) and surprisingly from there, the number of mutations reached 25 in January 2020 (**Figure S2**). We noticed that this number is increasing exponentially and reached 48 within December 2020. Form the monthly basis of sequence analysis, we observed that some of the samples from March, August, November, and December 2020 have 40 mutations whereas 35 mutations were observed in June, July and October (**Figure S1-S13**).

**Figure 1.**
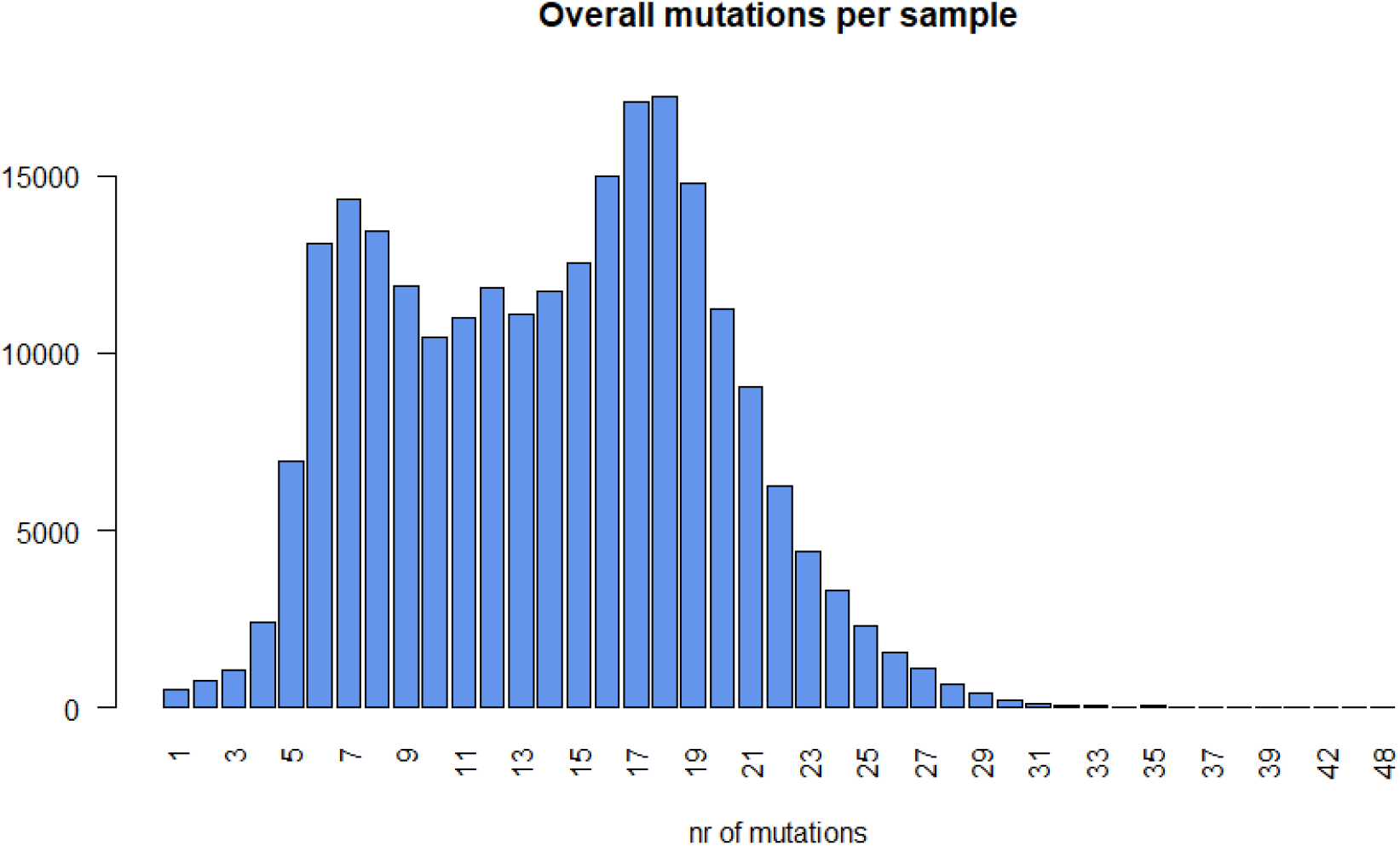
The number of mutational events occurred from December 2019 to December 2020 for all the 259044 SARS-CoV-2 genome samples. Y-axis represents the number of samples.

**Figure 2.**
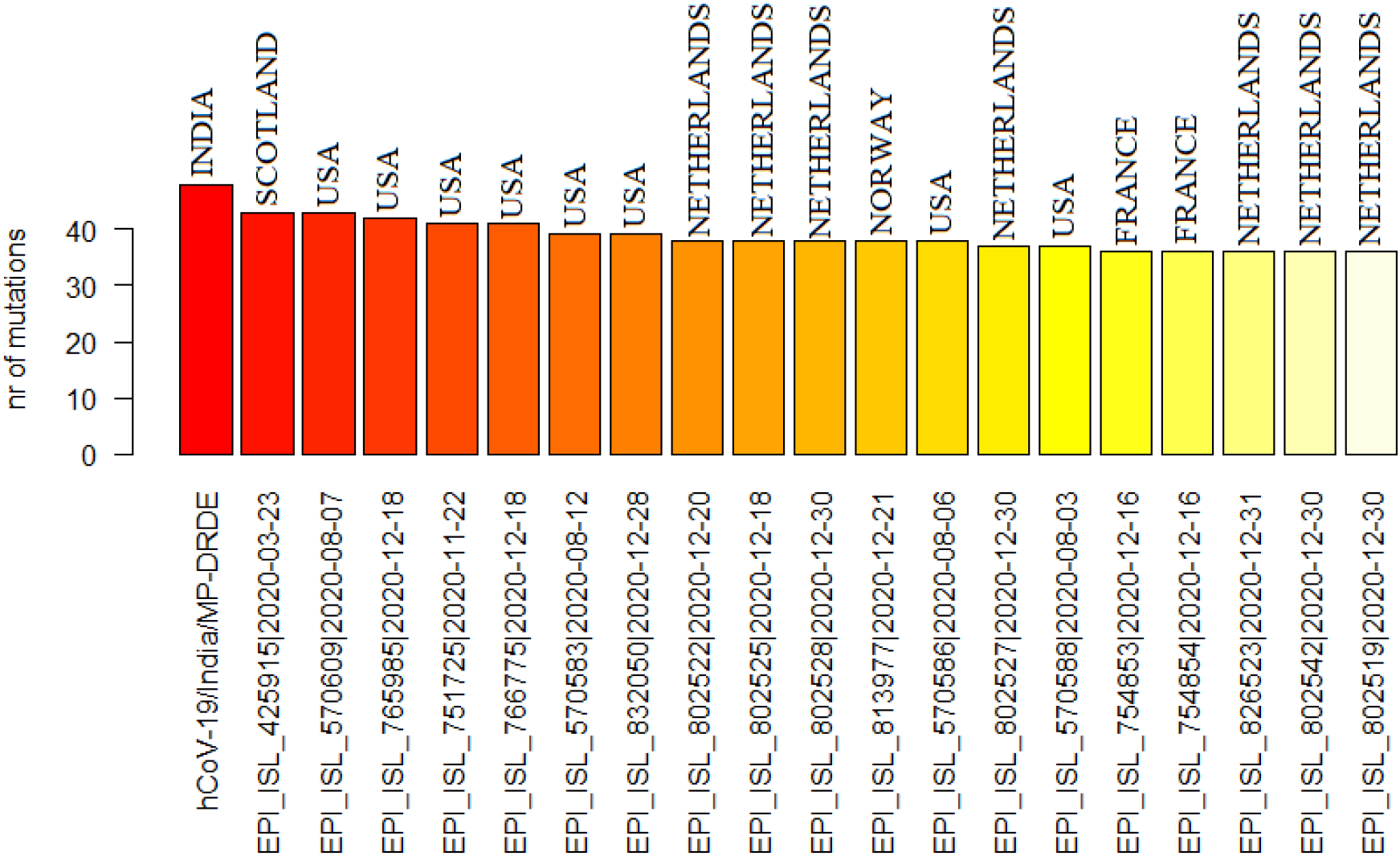
Distribution of the most mutated SARS-CoV-2 samples throughout the world from December 2019 to December2020.

### 2.2 Analysis of nature of SARS-CoV-2 mutations

The nature of each of 3334545 mutation had been analyzed and a high prevalence of a single nucleotide polymorphisms (SNPs) were identified worldwide over brief cases of deletion/insertion (indels). We observed a total of 1745775 missense mutations (52.35% of the total) where 1234456 nonsense mutations (37.02% of the total) were found to fall over the coding regions. Moreover, 337340 (10.11%), and 10220 (0.30 %) mutational events were identified in extragenic regions (5′ and 3′ untranslated region of the SARS-CoV-2 RNA sequence) and as deletion, respectively. Very small amount of stop codon creating SNP 4746 (0.14 %) were observed followed by the in-frame deletions, in-frame insertion, and insertion which account for 1122, 260 and 518 of all the investigated mutational cases (**Figure 3**). Similar type of mutational profile was also observed from monthly basis of sequence (December 2019 to December 2020) analysis which demonstrate that SNPs are the major mutational event followed by the silent SNP and intergenic mutations (**Figure S1-S13**), assuming a conserved molecular mechanism for SARA-CoV-2 mutational evolution.

**Figure 3.**
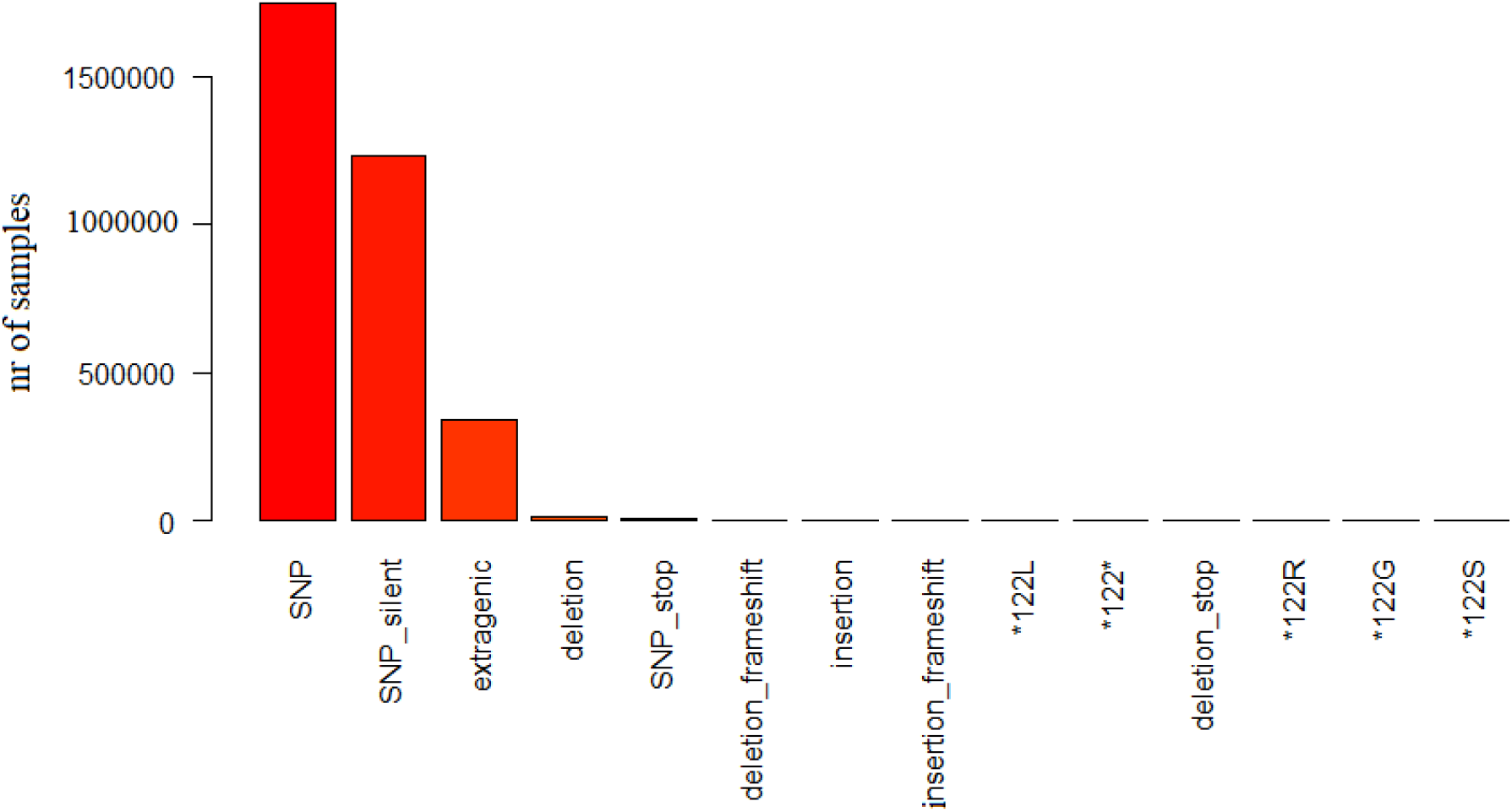
Most frequent classes of SARS-CoV-2 mutations throughout the world.

### 2.3 Analysis of SARS-CoV-2 mutations according to their type

To observe the pervasiveness of SNP transitions (purine to purine and pyrimidine to pyrimidine) and/or SNP trans-version (purine to pyrimidine and pyrimidine to purine) SARS-CoV-2 mutations were classified based on their types. Worldwide the most common transition event was noticed as C>T transition which accounts for 1756440 (52.67%) of all the observed 3334545 SARS-CoV-2 mutations. The second most common mutation type was marked as G>T trans-version with 486610 occurrences (14.59%) where A>G transition was noticed as third most common mutation type with 371334 (11.13%) cases. Transition (T>C, and G>A,) and trans-version (G>C, C>G, C>A, and A>T) were remarked as 4^th^, 5^th^, 6^th^, 8^th^, 9^th^, and 10^th^ common events of SARS-CoV-2 mutational evolution. A particular type of peculiar multi-nucleotide mutation (substitution of a GGG triplet with AAC) was observed as 7^th^ common mutational type worldwide with 67596 incidents. The deletion of the ATG codon is the most common indel for SARS-CoV-2 and was found as 16^th^ common mutational type with 1231 events (**Figure 4**). From our monthly basis mutation type analysis, we also noticed that the substitution of C with T is the most prominent alteration in every month from January 2020 to December 2020 worldwide (**Figure S2-S13**).

**Figure 4.**
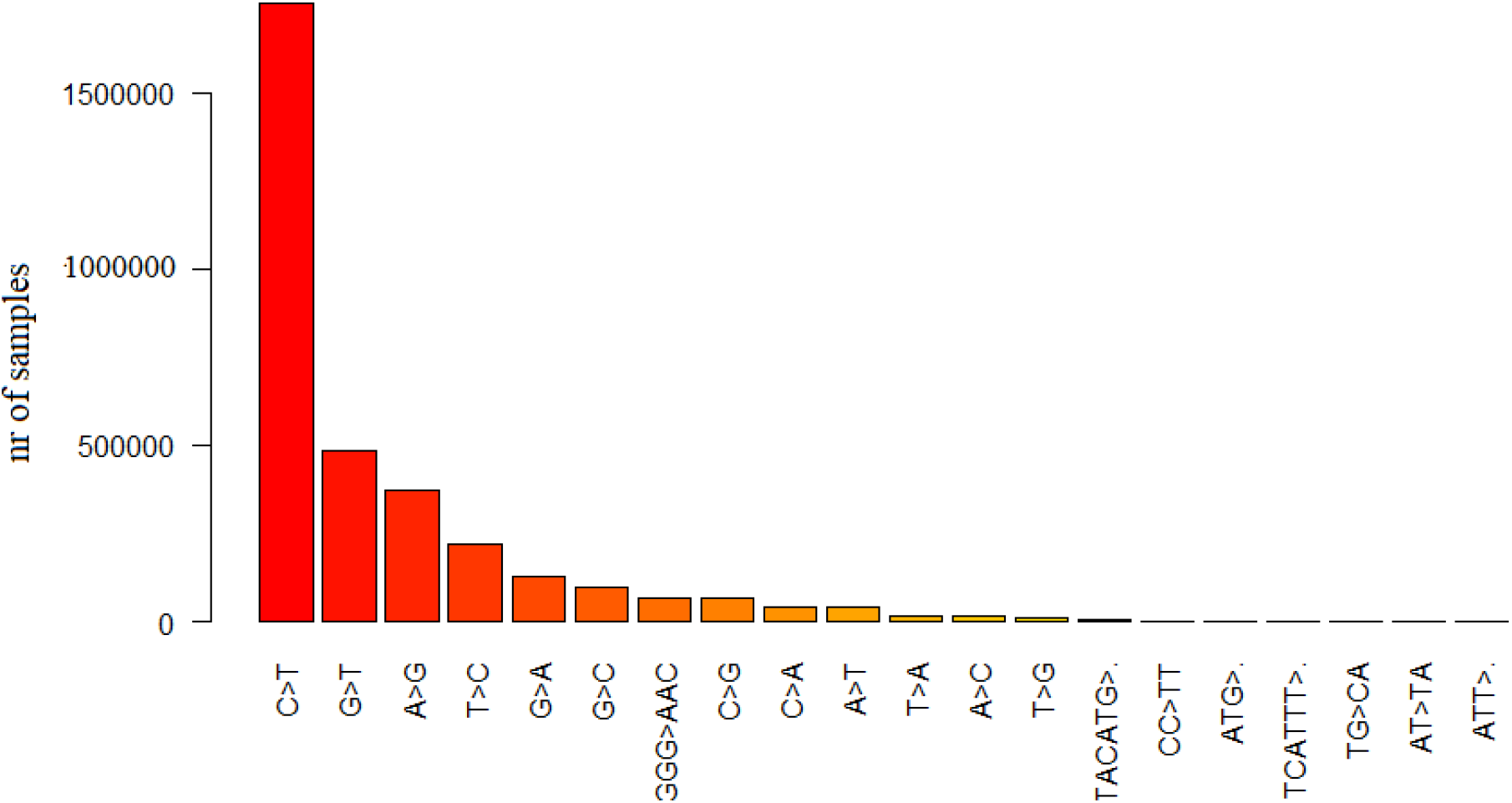
Most frequent types of SARS-CoV-2 mutations throughout the world.

### 2.4 Analysis of SARS-CoV-2 mutations in nucleotide and protein level

We investigated the effect of each genetic variation on the viral protein sequences. The most predominant mutations were observed in the 23403^th^ (A>G transition), 3037^th^ (C>T transition), 14408^th^ (C>T transition), and 241^th^ (C>T transition) nucleotide position of SARS-CoV-2 genome (**Figure 5**). The A23403G mutation causes a change from Aspartate to Glycine in protein position 614 (spike protein) which is responsible for the initial entry of the virus through the ACE2 receptor and associated with severity of COVID 19 ^19^. The C14408T mutation substitute Proline with Leucine in position 314 of non-structural protein 12 (NSP12), a RNA-dependent RNA polymerase (RdRp). Conversely, C3037T (F106F) mutation is found as a synonymous mutation in the region encoding NSP3, a viral predicted phosphoesterase whereas C241T mutations is falling over non-coding regions (5′ UTR) (**Figure 6**). Other common identified mutations are GGG28881AAC (RG203KR, in the Nucleocapsid protein N), C22227T (L93L, in the membrane protein M), G29645T (A222V, spike protein S), G21255C (A199A, in the NSP16), C28932T (V30L, in the ORF10), and T445C (A220V, in the N protein) (**Figure 5, 6**). Furthermore, monthly basis of sequence analysis revealed that D614G (S), F106F (NSP3), P314L (NSP12b), and 5′ UTR:241 mutations are in the top of the mutation analysis chart of every month from March 2020 to December 2020 (**Figure S4-S13**).

**Figure 5.**
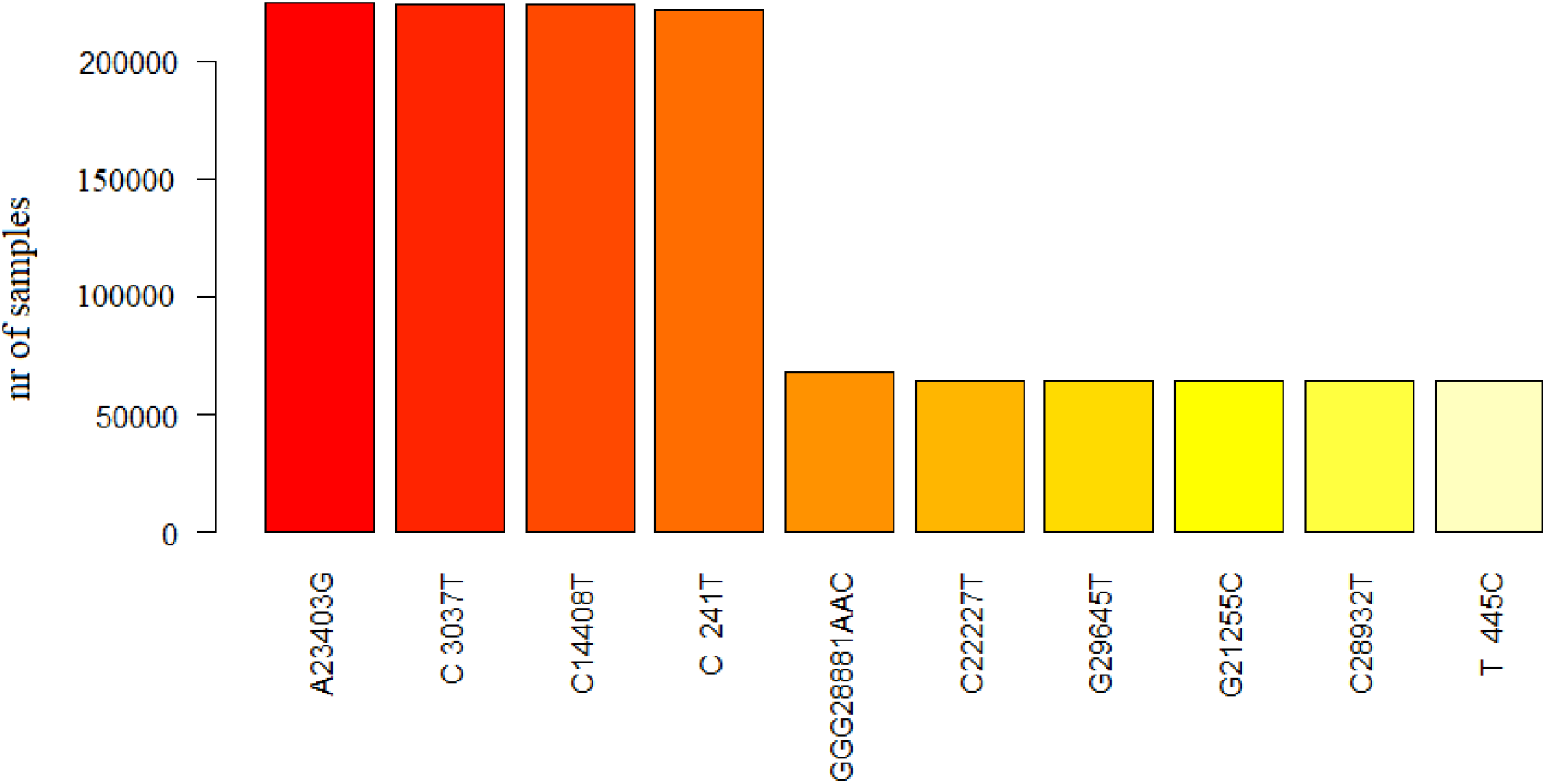
Worldwide distribution of most frequent SARS-CoV-2 mutational events (annotated a nucleotide coordinates over the reference genome).

**Figure 6.**
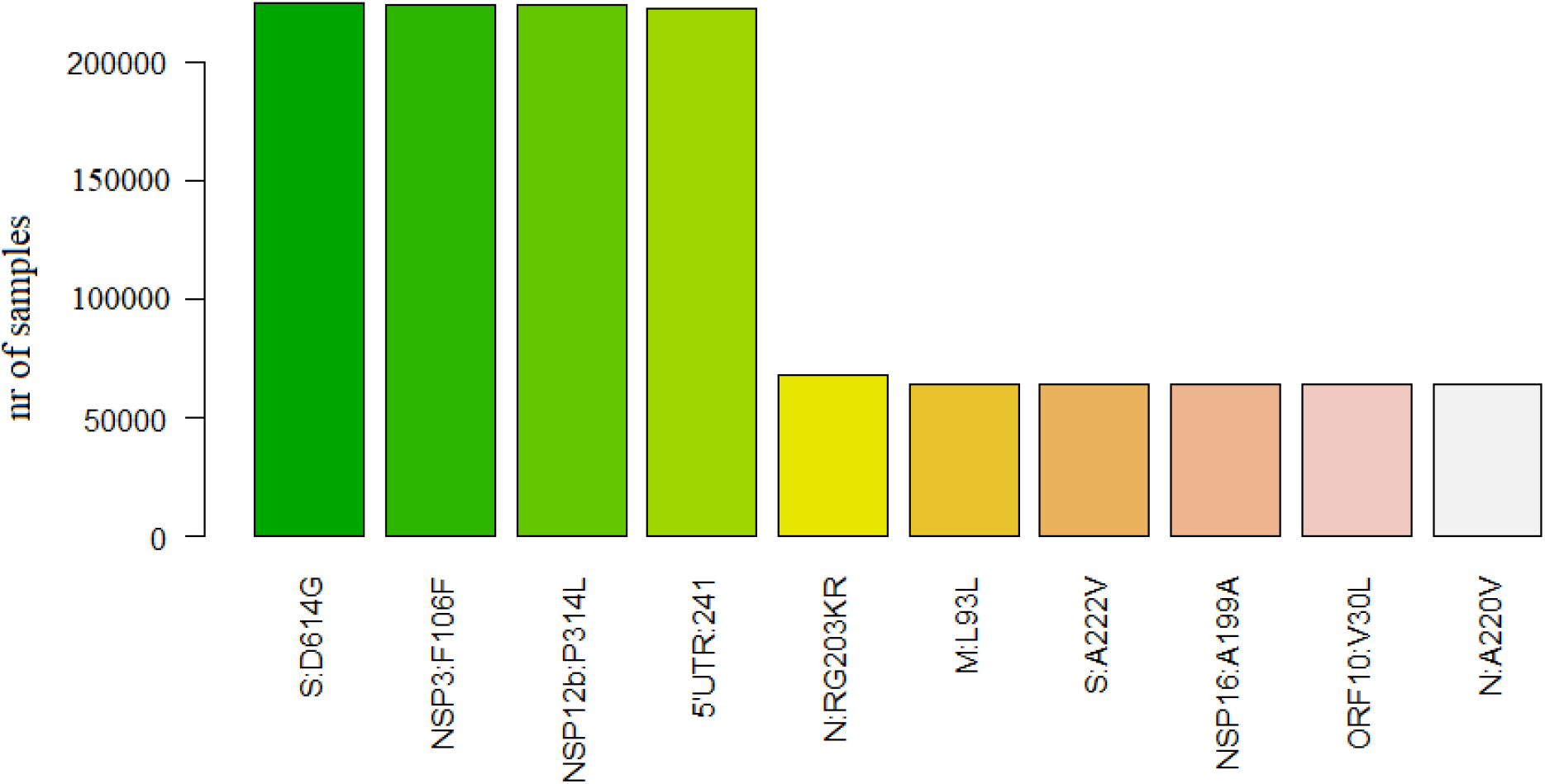
Worldwide distribution of most frequent SARS-CoV-2 mutational events (annotated a amino acid coordinates over the reference genome).

### 2.5 Prediction of mutation rate of future SARS-CoV-2 variants

Long Short Term Memory (LSTM) Network was used to build the mutation prediction model through the training and testing process of the COVID-19 patient’s sample. Our model can predict the mutation rate of 250 future variants for 98800 training samples at a time. To maintain the model size, we have added the immediately predicted samples at the bottom of the training samples and deducted the top 250 samples every time. Through this process we predicted th nucleotide mutation rate (%) for 2000 future variants. From this analysis, we observed that the nucleotide mutation rate between the 1^st^ (**Figure 7**) and 2000^th^ (**Figure 8**) future variants are highly deviated from each other. We noticed that the substitution of C with T is predominant in future variants and continuously increasing with the expansion of variants number and increased about 17% in case of 2000^th^ variant (56%) than 1^st^ (39%), suggesting a possible higher increment of C>T in future time. On the contrary, replacement of T with C was found to decrease about 7% in 2000^th^ variant (2%) than 1^st^ (9%), propounding a possible decrement of T>C in future time. In case of G>A and G>T substitutions, we perceived about 6% and 7% decrement respectively in 2000^th^ variant when compared to 1^st^ whereas 7% and 3% increment was respectively observed for A>G and A>T substitutions. We did not observe any noticeable alterations as T>G\A, C>G\A and A>T\C from 1^st^ to 2000^th^ future variants (**Figure 9**).

**Figure 7.**
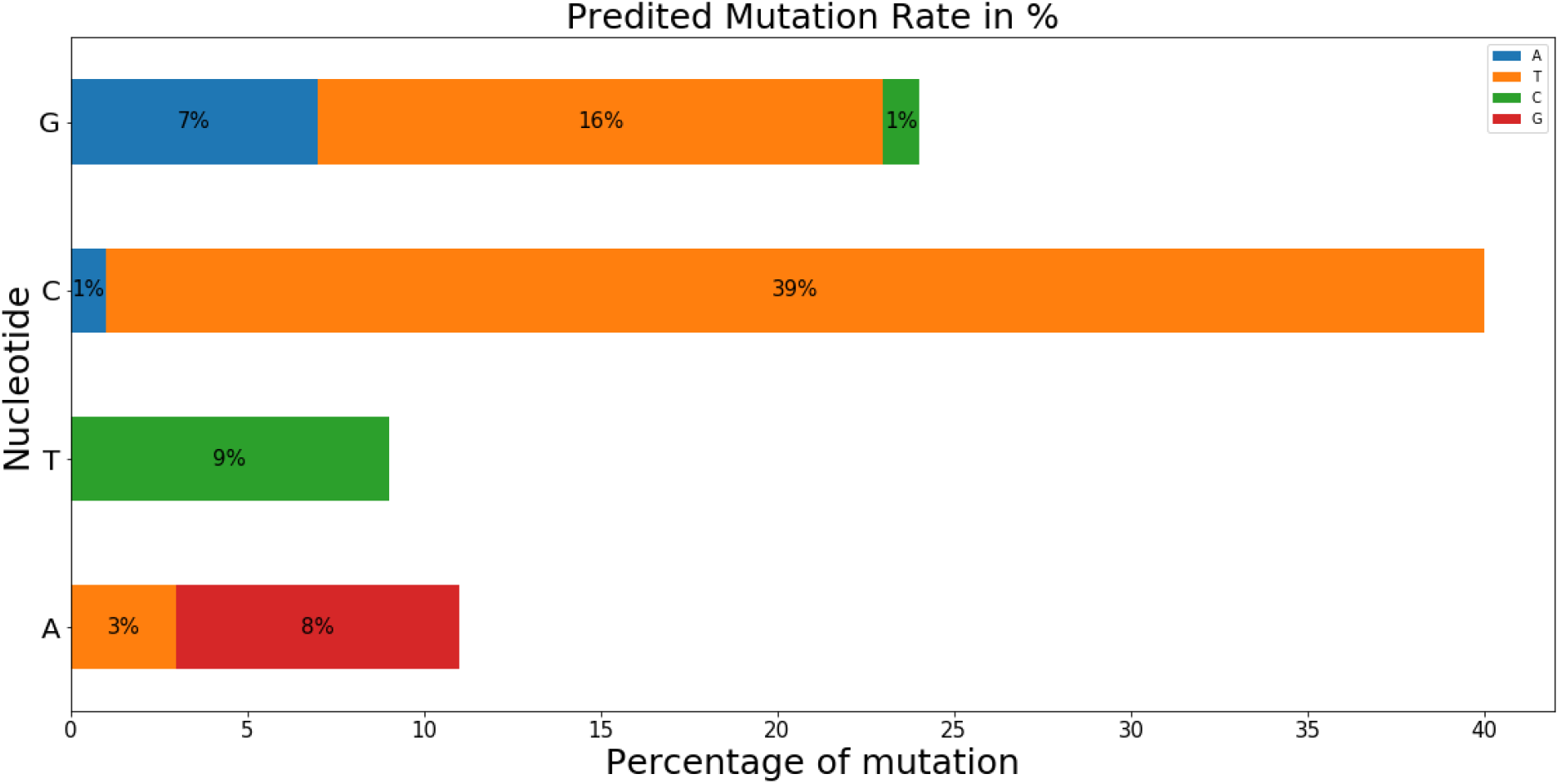
Predicted nucleotide mutation rate for 1^st^ future SARS-CoV-2 variant.

**Figure 8.**
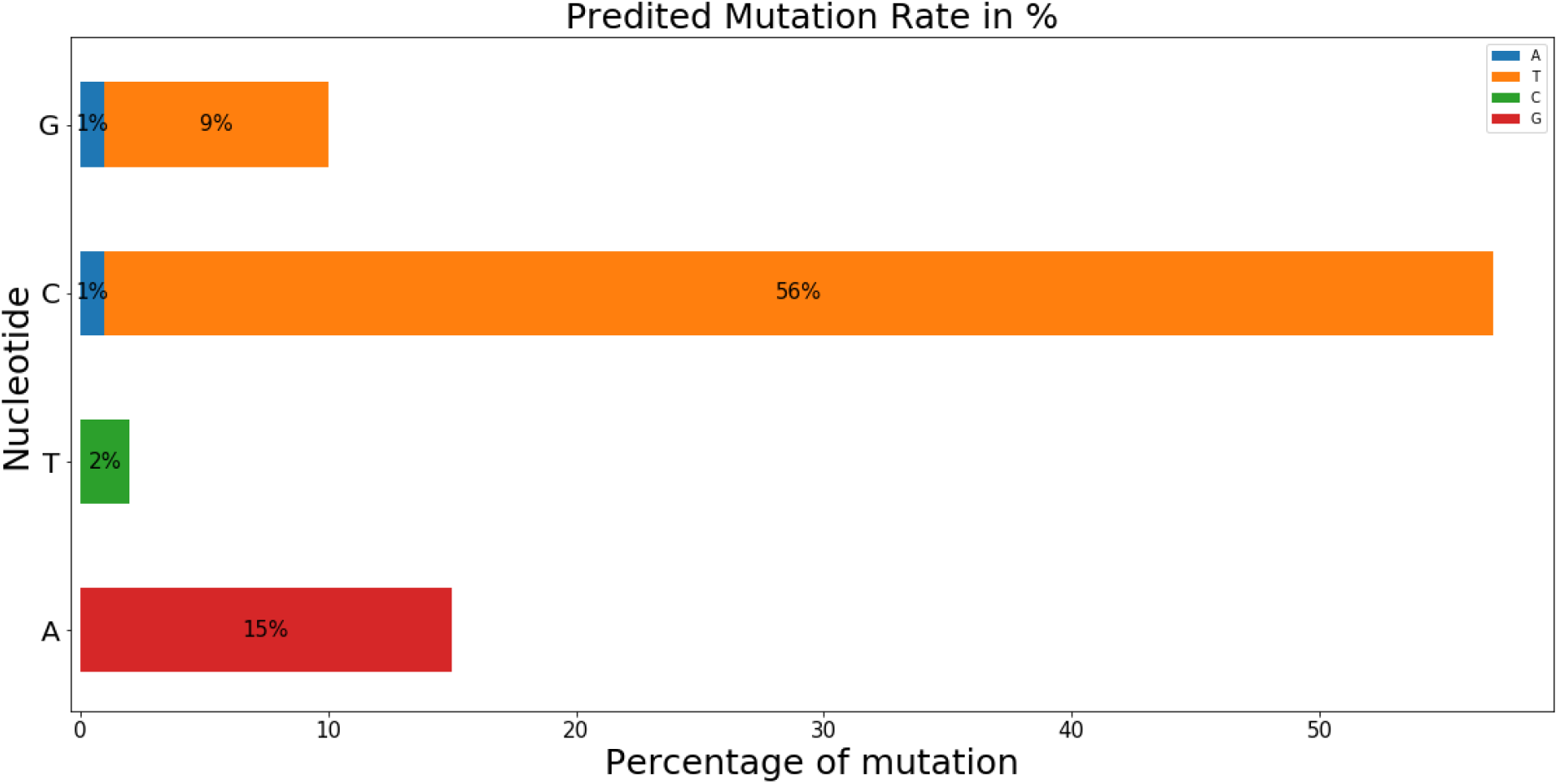
Predicted nucleotide mutation rate for 2000^th^ future SARS-CoV-2 variant.

**Figure 9.**
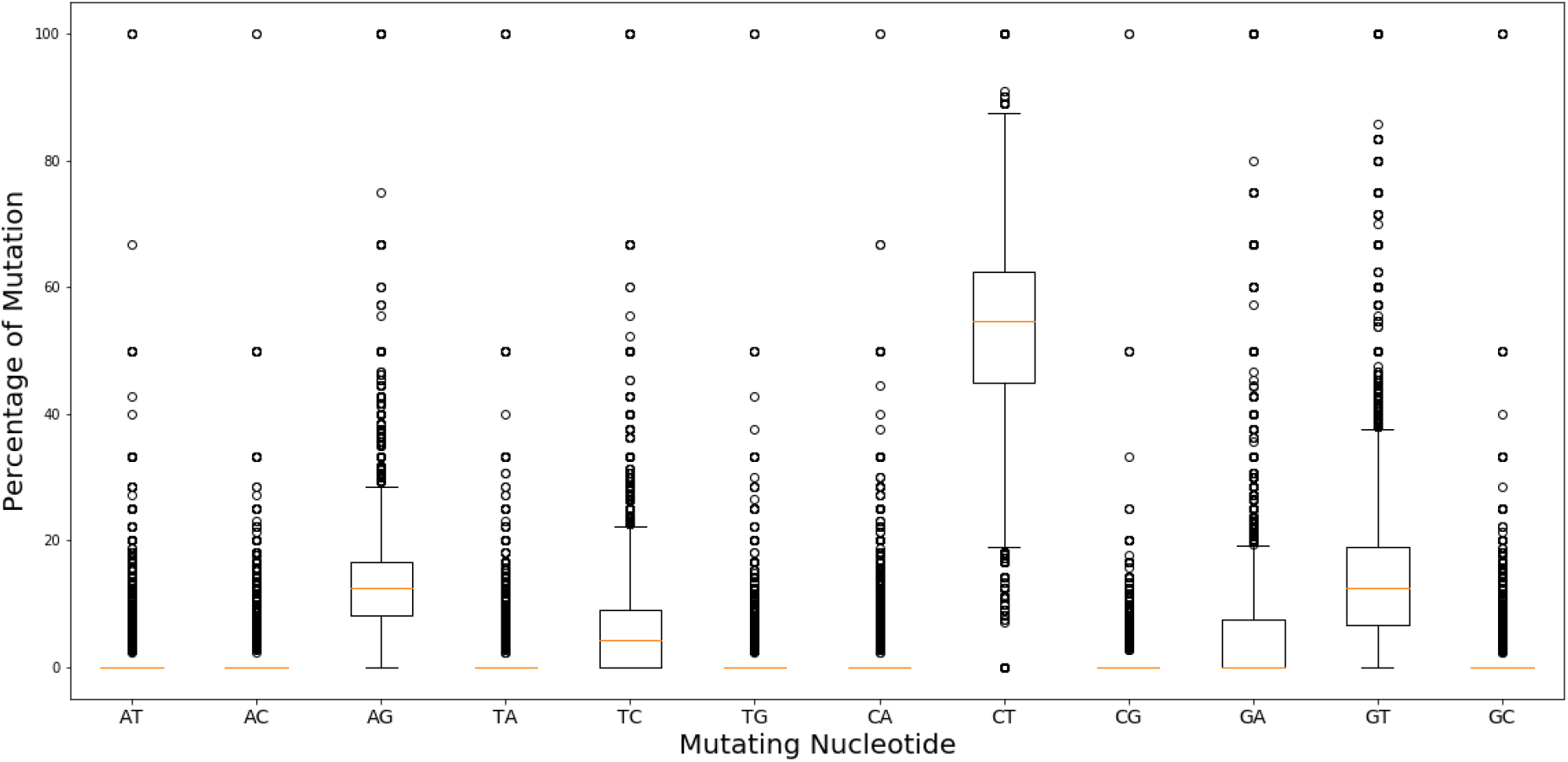
Overall predicted nucleotide mutation rate for 2000 future SARS-CoV-2 variants.

## 3. Discussion

Since SARS-CoV-2 is an RNA virus, it is evolving continuously in human populations over time and thus driving its massive worldwide spread. Due to the genetic diversity of the virus along with the patient’s genomic variations, the severity of COVID-19 greatly varies from patient to patient. A large proportion of patients either remain symptomless or show mild to moderate symptoms. Nevertheless, some patients who experience severe infection die after hospitalization ^20^. This difference in disease severity from one person to another is one of the mysteries events which is probably associated with the genetic variations of the virus. The genomic sequence data offers a great opportunities to study the molecular changes in the expanding viral population by providing new insights into the mode of spread, diversity during the pandemics and the dynamics of evolutions ^21^. The current study was intended to explore the genome wide accumulation of viral mutations at different time points to identify mutations which are occurring globally and to predict the mutation rate of the virus for future times through the artificial recurrent neural network (RNN) called Long Short Term Memory (LSTM).

Mutational profiles of the 259044 SARS-CoV-2 isolates from December 2019 to December 2020 recognize a total of 3334545 mutations with an average of 14.01 mutations per sample, confirms a relative high mutation rate of the virus. Each of the 17221, 17084, 14983, and 14801 samples were found to contain 18, 17, 16 and 19 mutations, respectively. Whereas 48 mutations were identified for 1 samples. The occurrence of such high amount of mutations in large number of samples indicating a faster molecular evolution of SARS-CoV-2 which probably responsible for making it more deadly in terms of time. Among the mostly mutated 20 samples, Indian sample had the maximum number of mutations of 48 followed by the samples (having up to 36 mutation) from Scotland, USA, Netherlands, Norway, and France. This high mutation rate could be the main reason behind the extremely deadly action of this virus in these particular regions specially USA and India. Alongside, we notice that the number of mutation is exponentially increased in every months from the emergence of SARS-CoV-2 till December 2020 (48 mutations) which is to suggest the virus holds its mutating nature and continuously evolving in the similar pattern. Analysis of the nature of SARS-CoV-2 mutations confirms a conserved molecular mechanism for SARA-CoV-2 mutational evolution as the missense mutations (52.35%) are the most prevalent mutational events in terms of long time, followed by the silent SNPs (37.02%) and extragenic SNPs (10.12%). Missense mutations are known to effect the protein structure, hence alter the stability even protein-protein interactions. Wide occurrence of missense mutations in the SARS-CoV-2 genome may change its proteins shape and stability that probably makes it more interactive with the host receptor protein ^19^. In our large scale study, previously reported D614G, and P314L missense mutations are also identified as the most prevalent mutation in the viral genome. The D614G mutation in spike protein is responsible for the initial entry of the virus through the hACE2 receptor and associated with severity of COVID 19. The P314L mutation in NSP12 is associated with the D614G mutation and may favor the SARS-CoV-2 by enhancing its transmission ability ^22^. Conversely, the nonsense mutations in the viral genome may alter the protein’s nature as they might cause complications with exon splicing enhancers (ESEs) which can change the mRNA processing ^23^. F106F mutation is found as predominantly occurring nonsense mutation in NSP3, suggesting a possible role in mRNA processing which might alter the nature of the viral protein. Moreover, extragenic SNP, specially SNPs in 5′UTR may affect the folding of the ssRNA and influence the transcription and replication rates of SARS-CoV-2 ^24^. With this term, the 5′ UTR:C241T mutation might be associated with the transcription and replication rates of SARS-CoV-2 as it is found to occur most prominently. Along with the previously found GGG>AAC mutation, our mutational type analysis identify some another multi-nucleotide mutations CC>TT, TG>CA, and AT>TA which are in the top 20 mutational type and should be monitored for the future as the GGG>AAC (R203K and G204R) reported to be associated with the insertion of a lysine in SR domain of N protein which might affect the phosphorylation ^25^. Besides D416G, F106F, P314L, and 5′ UTR:C241T, our large scale analysis also identify C22227T;L93L (membrane protein M), G29645T;A222V (spike protein S), G21255C;A199A (NSP16), C28932T;V30L (ORF10), and T445C;A220V (N protein) mutations which are in top 10 mutations found in our investigation and should get importance in evaluating their role in the efficiency of SARS-CoV-2 transmission. Whatever, since the SARS-CoV-2 continuously evolving, new mutations will surely emerge. Our Long Short Term Memory (LSTM) recurrent neural network based future mutation prediction model claim that the virus will continue to mutate in a high rate which might give evolutionary advantage to the virus as mutations occur in the similar fashion. Mutation rate analysis for future time for 2000 patients predicts a 17% increment of C>T with the expansion of patients number which suggesting a possible higher increment of C>T in future time. Furthermore, 7% and 3% increment is respectively observed for A>G and A>T substitutions. At the same time, decrement is also noticed for T>C (7%), G>A (6%), and G>T (7%) mutations in future time. The appearance of new mutations may affect the building of new therapies and even can impair the adaptation of current therapies to get rid of the new molecular structure of the coronavirus. The appearance of new mutations may increase the viral transmission. For example, we notice from our monthly sequence analysis that after the emergence of coronavirus in December 2019, the virus with a large number mutations (up to 30) were identified within 1 year in India, Scotland, USA, Netherlands, Norway, Israel, Italy, England and France, suggesting the fastest spread of SARS-CoV-2 subpopulation worldwide. Continuous tracking of mutations is the key to map the spread of the virus between individuals and throughout the world.

## 4. Conclusion

The COVID-19 has challenged the globe not just with regard to global health but also psychological and economic health. This challenge has been taken up in the scientific community and is investigating the virus and its pathogenicity along with clinical management. In this present study through integrated strategies of bioinformatics and deep neural learning, we explained the genetic variability of SARS-CoV-2 strains as well as the mutation rate of the virus in future. We also have explained the pattern of the identified mutations which showed a prevalence of single nucleotide polymorphism (SNPs) as a major mutational type worldwide. We highlighted the viral strains containing highest number of mutations. Most frequently occurring mutations in protein level was also explained along with multi-nucleotide mutations. Based on the LSTM-RNN model, we predicted the nucleotide mutation rate for 2000 future variants which exhibited that the virus will continue to evolve in a similar pattern, suggesting a conserved molecular evolution of the virus. To the best our knowledge, this study will facilitate the tracking of mutations and help to map the spread of the virus worldwide. Extensive studies will be required to explore whether the identified mutations can exert any effect on the COVID-19 severity.

## 5. Methods

### 5.1 Genomic data retrieval of SARS-CoV-2

To investigate the genetic variations of SARS-CoV-2 virus genomes, we retrieved 259044 complete genome sequences submitted from 01 December 2019 to 31 December 2020 across all countries from the Global Influenza Data Sharing Initiative (GISAID) database ^26^. The full-length sequences (>29000bp), as well as high nucleotide coverage (<1% Ns; <0.05% unique amino acid mutations), were considered for retrieving the SARS-CoV-2 genome sequences. Genomes associated with human infection were taken and the poor coverage (>5 percent Ns) genomes were excluded from the list. The reference genome sequence (NC_045512.2) of SARS-CoV-2 ^27^ containing 29,903bp in length was downloaded in FASTA format from National Center for Biotechnological Information (NCBI) database.

### 5.2 Genomic variability analysis of SARS-CoV-2

A GFF3 annotation file associated to the reference was used to show the genomic sequences for all protein sequences of SARS-CoV-2. The ORF1 polyprotein was divided into its Non-structural proteins (NSPs) constituent such as NSP12. In the genome annotation, viral RNA-dependent RNA polymerase encoding NSP12 was considered as two regions called NSP12a and NSP12b. Alignment of total 259044 SARS-CoV-2 genome sequences was done by using NUCMER v3.1 algorithm ^28^ where NC_045512.2 was considered as reference sequence. An R script was used to convert the output of the alignment to an annotated variant list that contain all the mutational events in nucleotide and protein level ^29^. The annotated variant list was then loaded through an in-house R script and checked whether there is any IUPAC code other than A, C, G, and T. If there is a different code, the list is fixed by removing it. Finally, the reference sequence was loaded along with the GFF3 annotation file of the reference sequence. The NUCMER object was then categorized according to SNPs, insertions and deletions. All the SNPs were then merged together in a separate file. Similarly, all the insertions and deletions were separated. Afterwards, we again merged the NUCMER object according to neighboring events (SNPs, insertions and deletions). Changes in query protein sequence according to variants were observed over the GFF3 annotation file of the reference sequence.

### 5.3 Deep neural learning and future mutation prediction

A dataset having all the nucleotide mutation data from 01 December 2019 to 31 December 2020 was prepared and processed to predict future mutations based on machine learning approach. In this regard, each sample with single or multiple mutations was separated and labeled. For each mutation (A>T\C\G, T>A\C\G, C>A\T\G, and G>A\T\C), the mutation rate (%) was calculated for each sample. Afterwards, the dataset that contained percentage based mutation data was used to select around 100000 samples randomly to run the model properly within our limited resources. The selected data were later divided to 80/20% as train and test sets. The train set was scaled by MinMaxScaler() function, a function of Scikit-learn machine learning library, and a time series generator was defined for the prediction of future mutations ^30^.

An artificial recurrent neural network (RNN) called Long Short Term Memory (LSTM) Network was used to build the mutation prediction model. The model was trained with a TimeseriessGenerator, a tool of keras API used for automatically transform a univariate or multivariate time series dataset into a supervised learning problem, and compared against the Test set. The input layer of the model got the prepared set of training data with 200 neurons. Then it has been through a dense layer of 200 neurons with relu activation layer. After that 0.15 dropout has been used. A dense of 12 neurons has been used as an output layer. The model was trained in 100 epochs. Adam optimized and mse (Mean Squared Error) loss function was used to train the model. Finally, the mutation rate (%) was predicted for the future 2000 SARS-CoV-2 variants.

## 6. Data availability

The datasets generated during and/or analyzed during the current study are available from the corresponding author on reasonable request.

## 8. Acknowledgments

The authors acknowledge the Department of Biotechnology and Genetic Engineering (BGE), Noakhali Science and Technology University and Computational Biology and Chemistry lab (CBC) for providing the research facilities.

## 9. Author contribution

Md. Shahadat Hossain, Md. Mizanur Rahaman developed the hypothesis. A. Q. M. Sala Uddin Pathan, Md. Nur Islam, Md. Shahadat Hossain, Mahafujul Islam Quadery Tonmoy, Mahmudul Islam Rakib, Md. Adnan Munim performed the study. Mahafujul Islam Quadery Tonmoy, A. Q. M. Sala Uddin Pathan, Atqiya Fariha, Otun Saha wrote the manuscript. Md. Shahadat Hossain, A. Q. M. Sala Uddin Pathan, Hasan Al Reza, Maitreyee Roy, Newaz Mohammed Bahadur, Md. Mizanur Rahaman reviewed the manuscript. All authors approved the manuscript.

## References

1 Zhu, N. et al. A novel coronavirus from patients with pneumonia in China, 2019. New England journal of medicine (2020).

2 Wu, F. et al. A new coronavirus associated with human respiratory disease in China. Nature 579, 265–269 (2020).

3 Dowd, J. B. et al. Demographic science aids in understanding the spread and fatality rates of COVID-19. Proceedings of the National Academy of Sciences 117, 9696–9698 (2020).

4 Geoghegan, J. L., Senior, A. M., Di Giallonardo, F. & Holmes, E. C. Virological factors that increase the transmissibility of emerging human viruses. Proceedings of the National Academy of Sciences 113, 4170–4175 (2016).

5 Pachetti, M. et al. Emerging SARS-CoV-2 mutation hot spots include a novel RNA-dependent-RNA polymerase variant. Journal of translational medicine 18, 1–9 (2020).

6 H, L. et al. Mutations: Types and Causes. Four edn, (2000).

7 Sanjuán, R., Nebot, M. R., Chirico, N., Mansky, L. M. & Belshaw, R. Viral mutation rates. Journal of virology 84, 9733–9748 (2010).

8 Vignuzzi, M., Stone, J. K., Arnold, J. J., Cameron, C. E. & Andino, R. Quasispecies diversity determines pathogenesis through cooperative interactions in a viral population. Nature 439, 344–348 (2006).

9 Chiara, M., Horner, D. S., Gissi, C. & Pesole, G. Comparative genomics suggests limited variability and similar evolutionary patterns between major clades of SARS-Cov-2. BioRxiv (2020).

10 Armijos-Jaramillo, V., Yeager, J., Muslin, C. & Perez-Castillo, Y. SARS-CoV-2, an evolutionary perspective of interaction with human ACE2 reveals undiscovered amino acids necessary for complex stability. Evolutionary Applications 13, 2168–2178 (2020).

11 Su, Y. C. et al. Discovery and genomic characterization of a 382-nucleotide deletion in ORF7b and ORF8 during the early evolution of SARS-CoV-2. MBio 11 (2020).

12 Gong, Y.-N. et al. SARS-CoV-2 genomic surveillance in Taiwan revealed novel ORF8-deletion mutant and clade possibly associated with infections in Middle East. Emerging microbes & infections 9, 1457–1466 (2020).

13 Benvenuto, D. et al. The 2019-new coronavirus epidemic: evidence for virus evolution. Journal of medical virology 92, 455–459 (2020).

14 Rubino, S., Kelvin, N., Bermejo-Martin, J. F. & Kelvin, D. As COVID-19 cases, deaths and fatality rates surge in Italy, underlying causes require investigation. The Journal of Infection in Developing Countries 14, 265–267 (2020).

15 Coppée, F., Lechien, J., Declèves, A.-E., Tafforeau, L. & Saussez, S. Severe acute respiratory syndrome coronavirus 2: virus mutations in specific European populations. New microbes and new infections 36, 100696 (2020).

16 Saha, O., Hossain, M. S. & Rahaman, M. M. Genomic exploration light on multiple origin with potential parsimony-informative sites of the severe acute respiratory syndrome coronavirus 2 in Bangladesh. Gene reports 21, 100951 (2020).

17 Saha, O. et al. Temporal landscape of mutation accumulation in SARS-CoV-2 genomes from Bangladesh: possible implications from the ongoing outbreak in Bangladesh. bioRxiv (2020).

18 Ojosnegros, S. & Beerenwinkel, N. Models of RNA virus evolution and their roles in vaccine design. Immunome research 6, 1–14 (2010).

19 Korber, B. et al. Tracking changes in SARS-CoV-2 Spike: evidence that D614G increases infectivity of the COVID-19 virus. Cell 182, 812-827.e819 (2020).

20 Callaway, E., Ledford, H. & Mallapaty, S. Six months of coronavirus: the mysteries scientists are still racing to solve. Nature 583, 178–179 (2020).

21 Khailany, R. A., Safdar, M. & Ozaslan, M. Genomic characterization of a novel SARS-CoV-2. Gene reports 19, 100682 (2020).

22 Wang, R. et al. Analysis of SARS-CoV-2 mutations in the United States suggests presence of four substrains and novel variants. Communications biology 4, 1–14 (2021).

23 Dickson, E. T. & Hyman, P. Brenner’s Encyclopedia of Genetics. Second edn, (Elsevier, 2013).

24 Kim, D. et al. The architecture of SARS-CoV-2 transcriptome. Cell 181, 914-921.e910 (2020).

25 Ayub, M. I. Reporting two SARS-CoV-2 strains based on a unique trinucleotide-bloc mutation and their potential pathogenic difference. (2020).

26 Shu, Y. & McCauley, J. GISAID: Global initiative on sharing all influenza data–from vision to reality. Eurosurveillance 22, 30494 (2017).

27 Gorbalenya, A. E. et al. Coronaviridae Study Group of the International Committee on Taxonomy of Viruses. The species severe acute respiratory syndrome-related coronavirus: classifying 2019-nCoV and naming it SARS-CoV-2. Nat. Microbiol 5, 536–544 (2020).

28 Delcher, A. L., Phillippy, A., Carlton, J. & Salzberg, S. L. Fast algorithms for large-scale genome alignment and comparison. Nucleic acids research 30, 2478–2483 (2002).

29 Mercatelli, D. & Giorgi, F. M. Geographic and genomic distribution of SARS-CoV-2 mutations. Frontiers in microbiology 11, 1800 (2020).

30 Pathan, R. K., Biswas, M. & Khandaker, M. U. Time series prediction of COVID-19 by mutation rate analysis using recurrent neural network-based LSTM model. Chaos, Solitons & Fractals 138, 110018 (2020).

