## Supplementary material for "Genome-wide identification and prediction of SARS-CoV-2 mutations show an abundance of variants: Integrated study of bioinformatics and deep neural learning": Figure S1-13

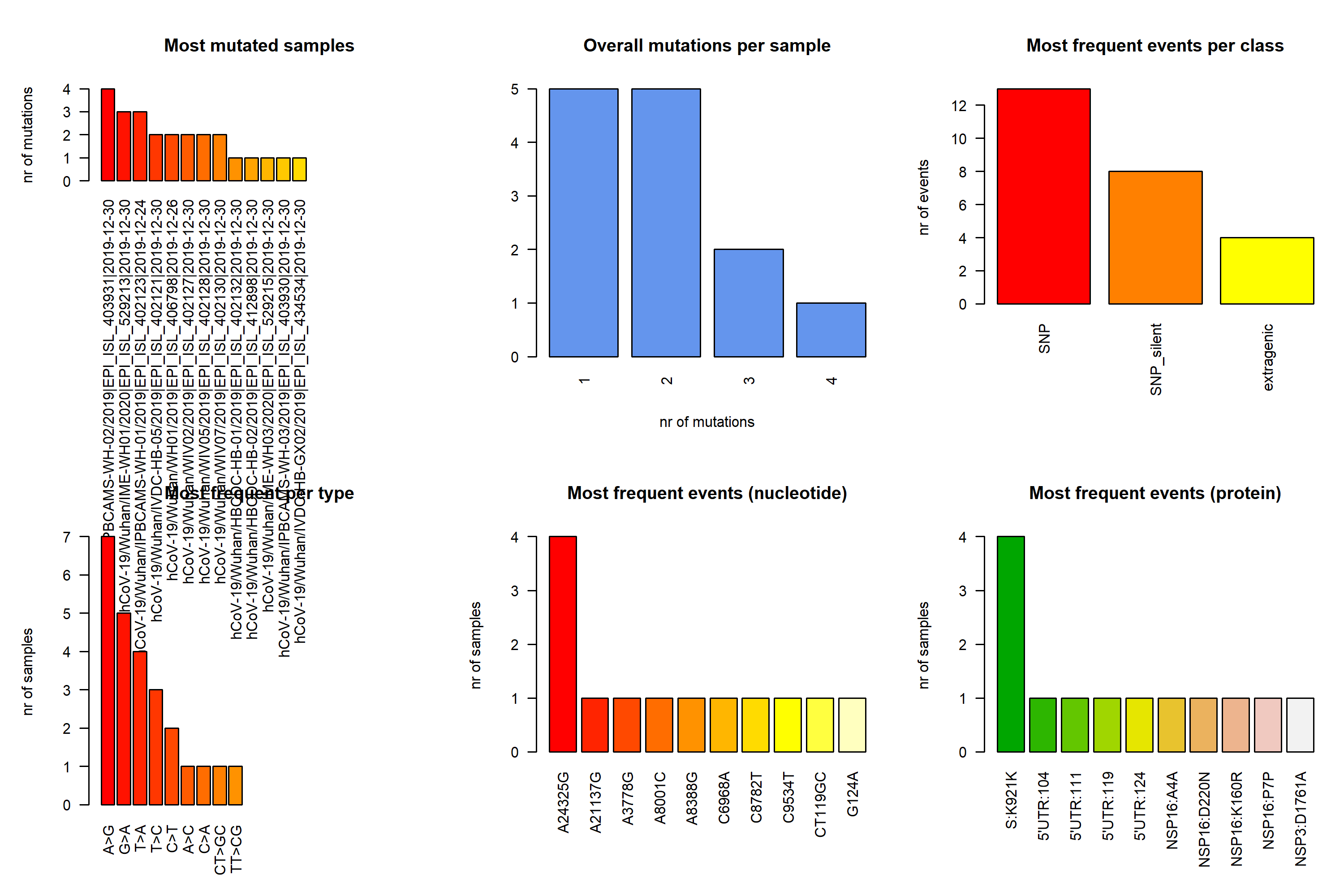

**Figure S1:** Mutational analysis of SARS-CoV-2 genome based on the sequencing data from December 2019.

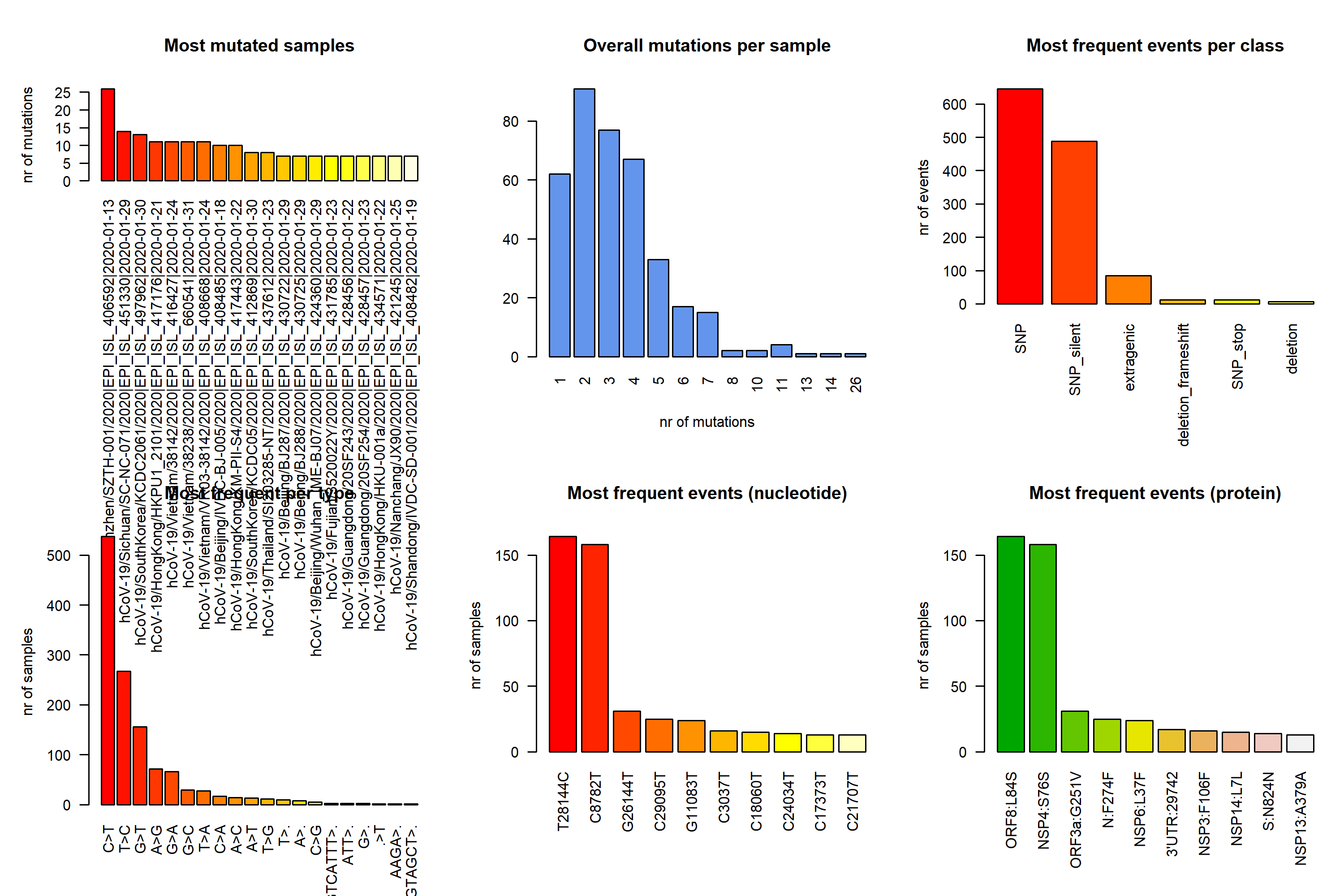

**Figure S2:** Mutational analysis of SARS-CoV-2 genome based on the sequencing data January 2020.

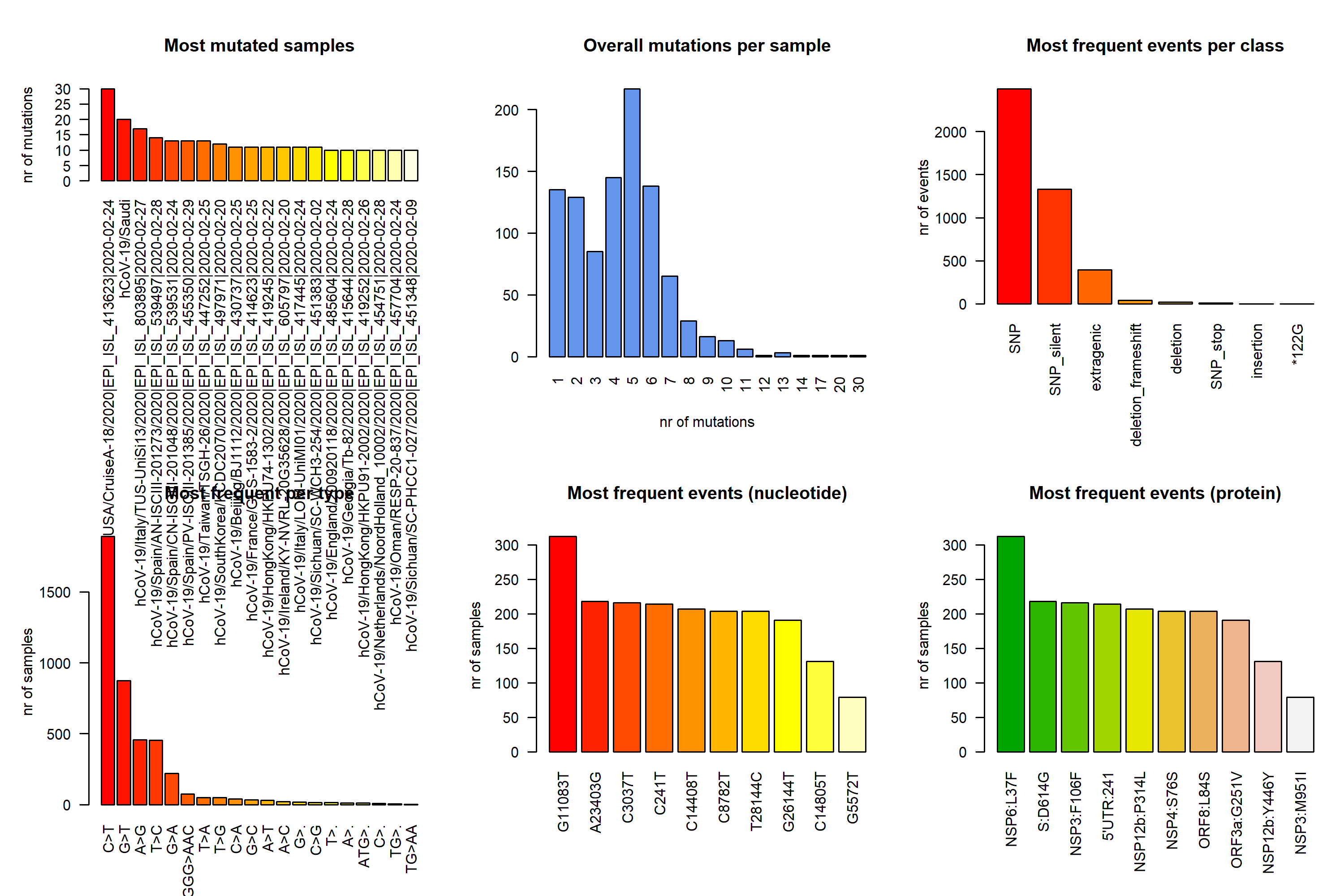

**Figure S3:** Mutational analysis of SARS-CoV-2 genome based on the sequencing data February 2020.

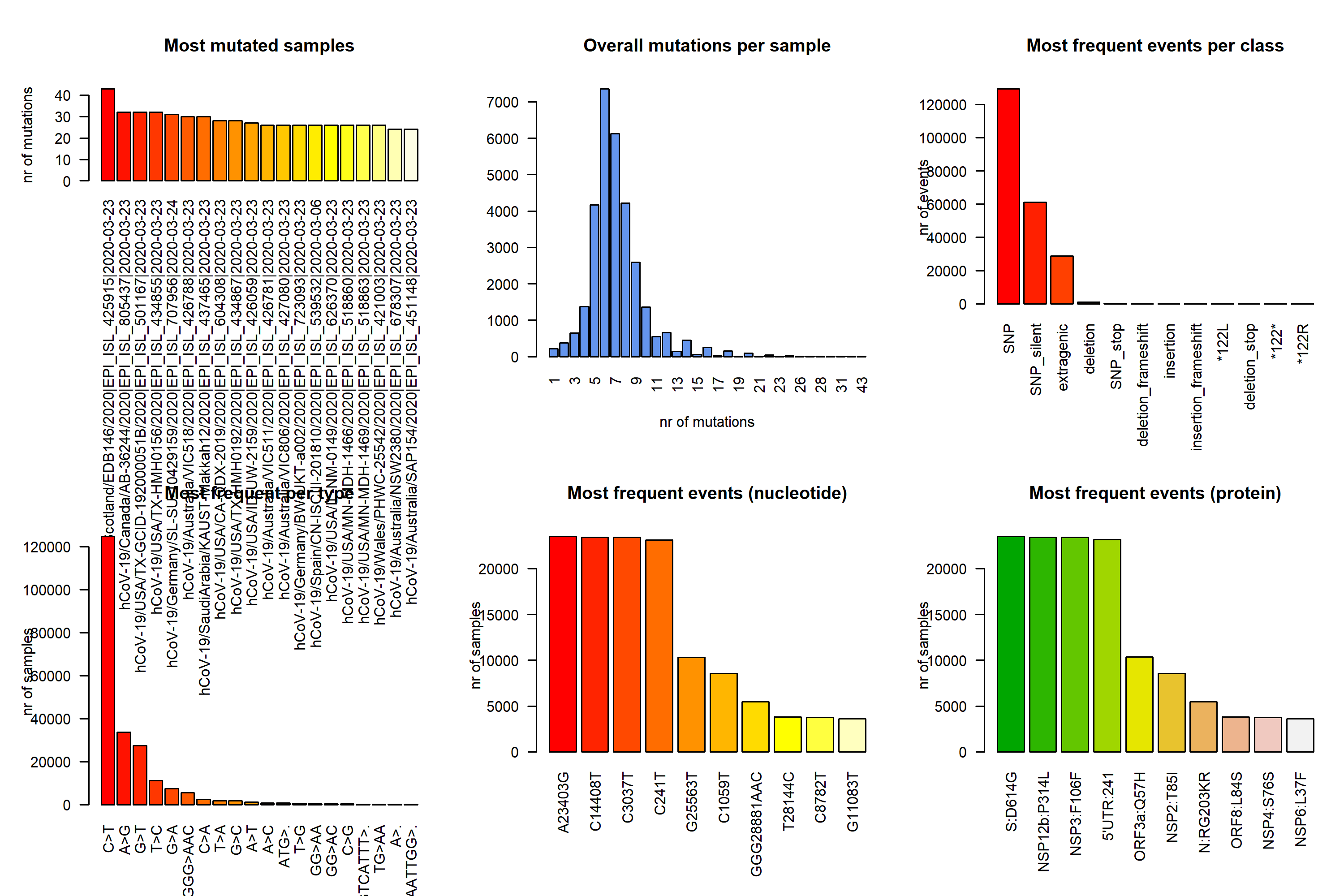

**Figure S4:** Mutational analysis of SARS-CoV-2 genome based on the sequencing data March 2020.

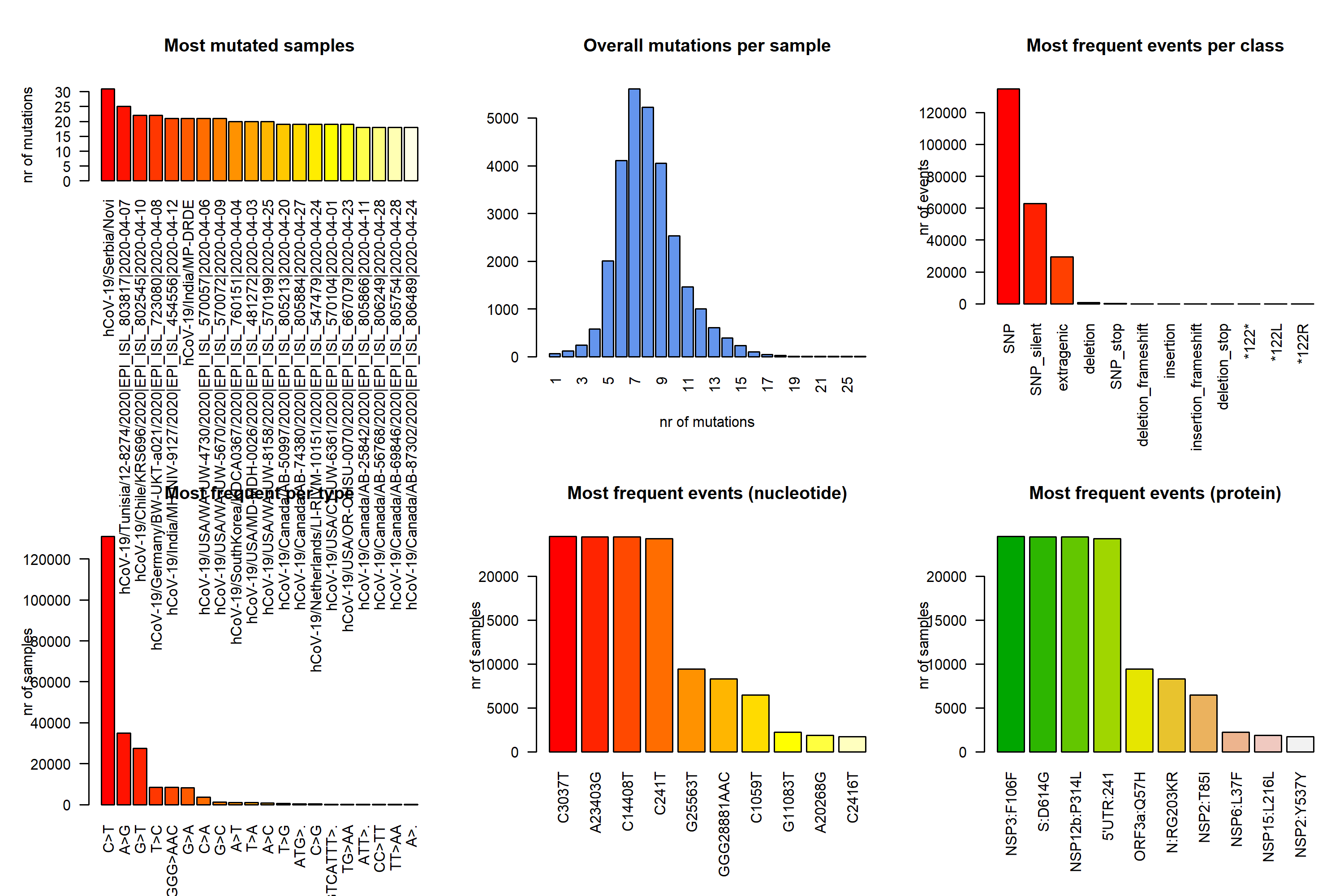

**Figure S5:** Mutational analysis of SARS-CoV-2 genome based on the sequencing data April 2020.

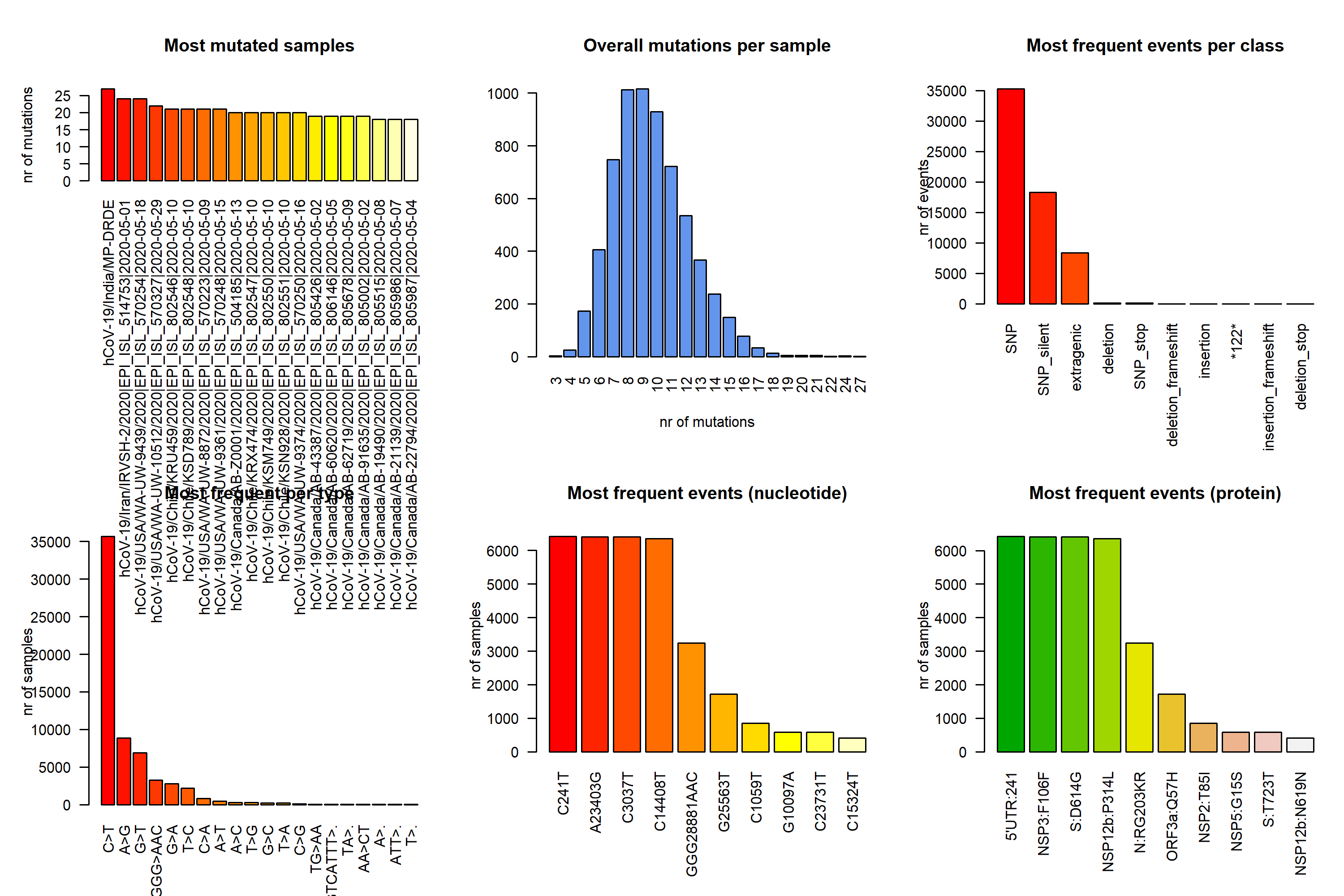

**Figure S6:** Mutational analysis of SARS-CoV-2 genome based on the sequencing data May 2020.

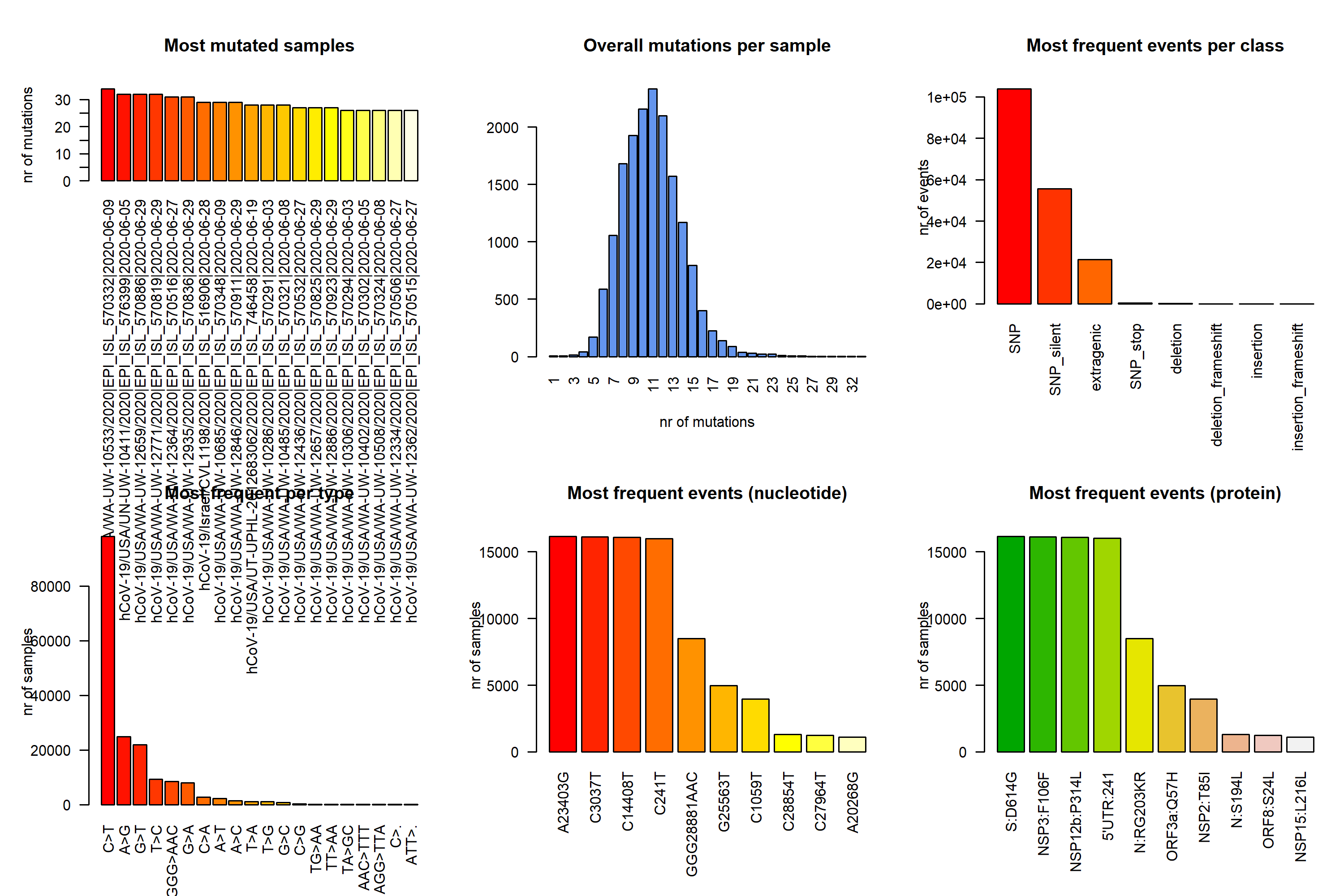

**Figure S7:** Mutational analysis of SARS-CoV-2 genome based on the sequencing data June 2020.

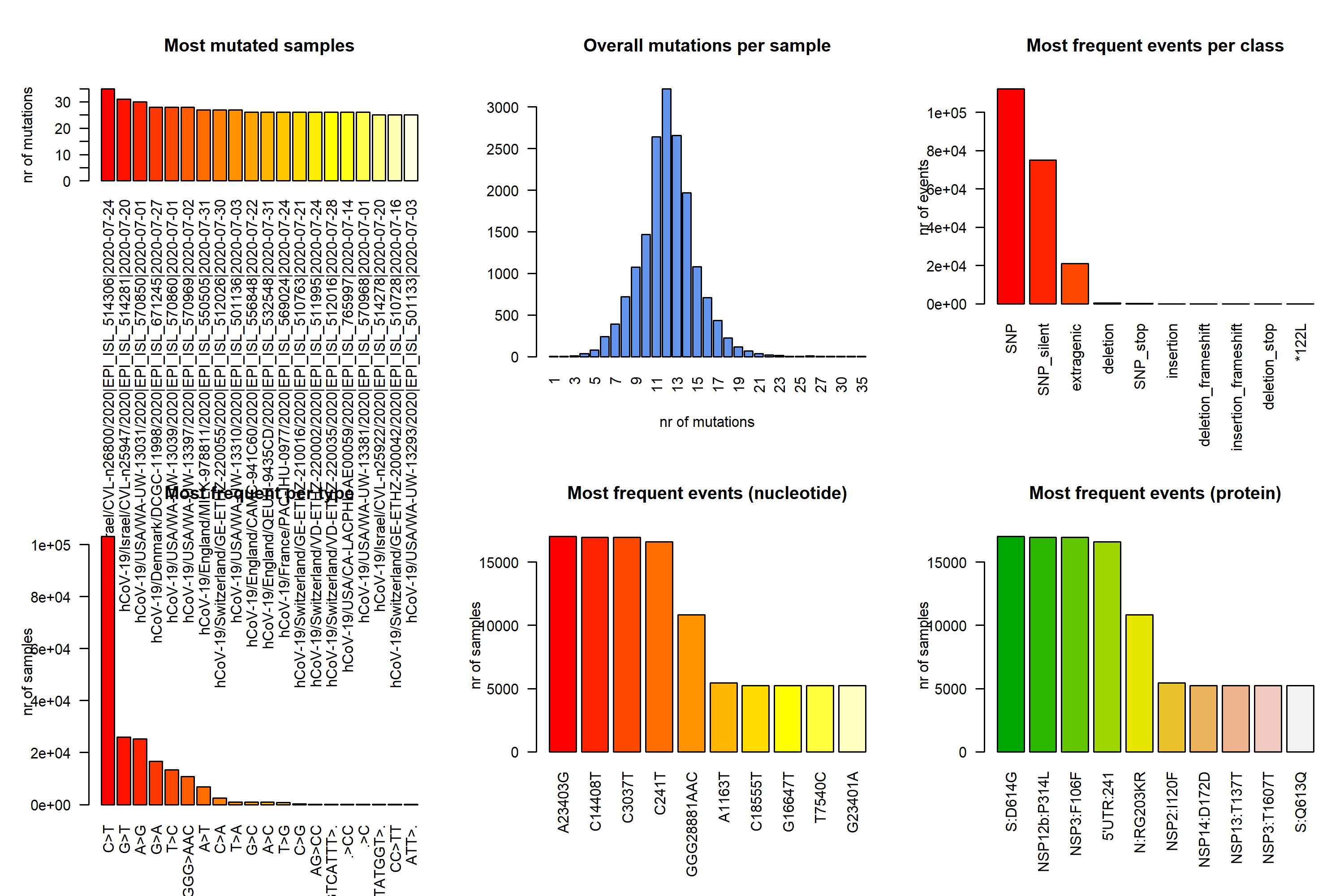

**Figure S8:** Mutational analysis of SARS-CoV-2 genome based on the sequencing data July 2020.

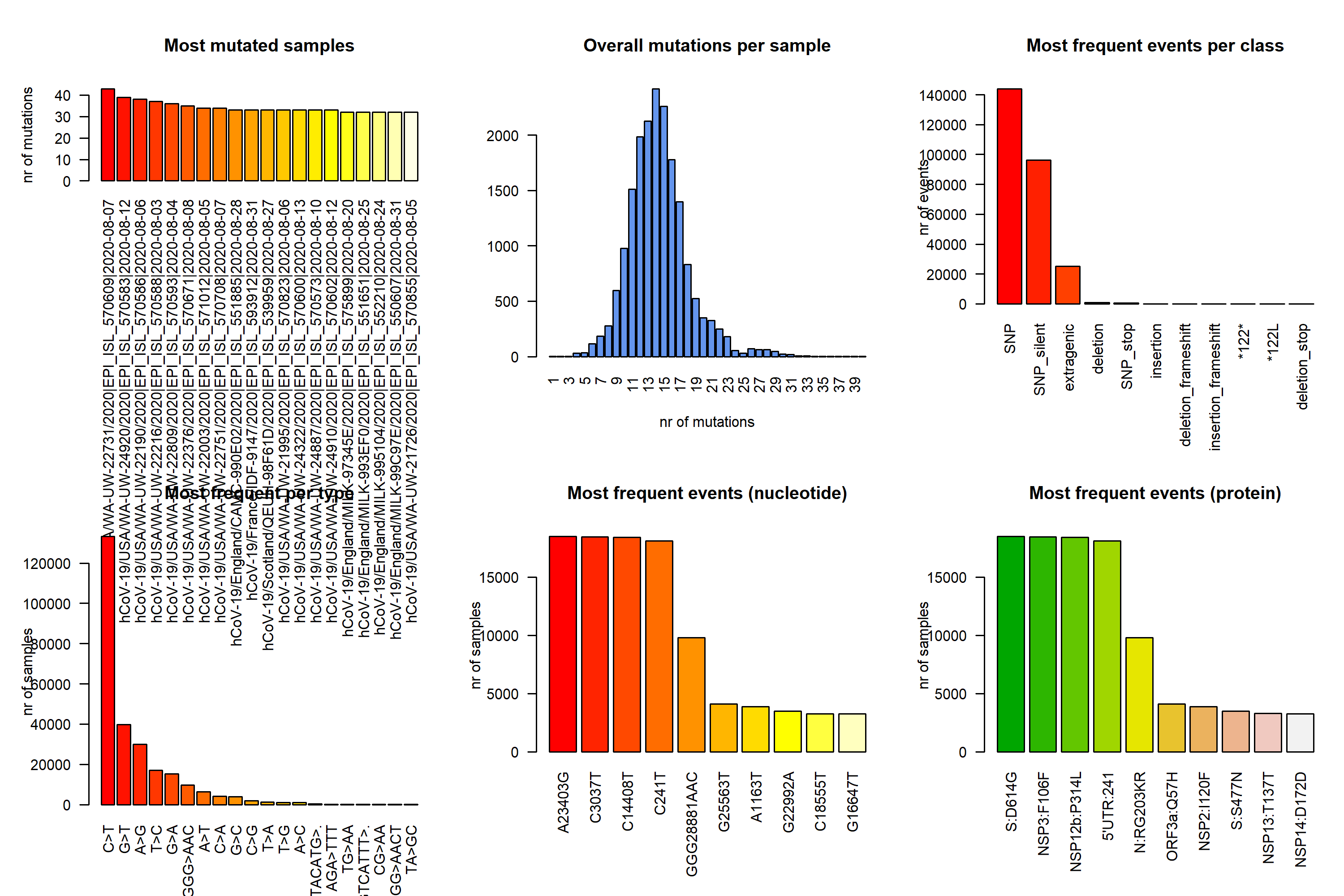

**Figure S9:** Mutational analysis of SARS-CoV-2 genome based on the sequencing data August 2020.

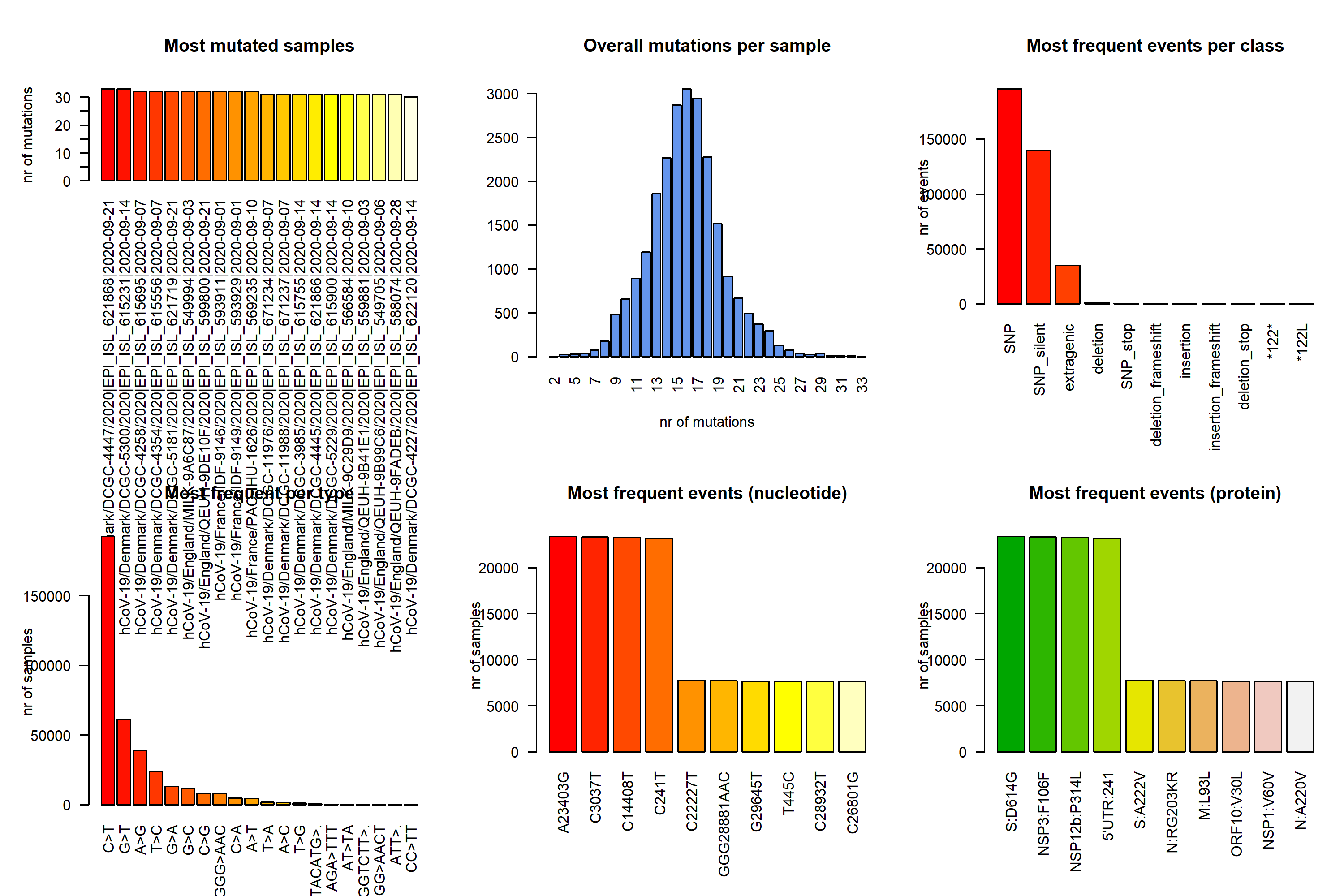

**Figure S10:** Mutational analysis of SARS-CoV-2 genome based on the sequencing data September 2020.

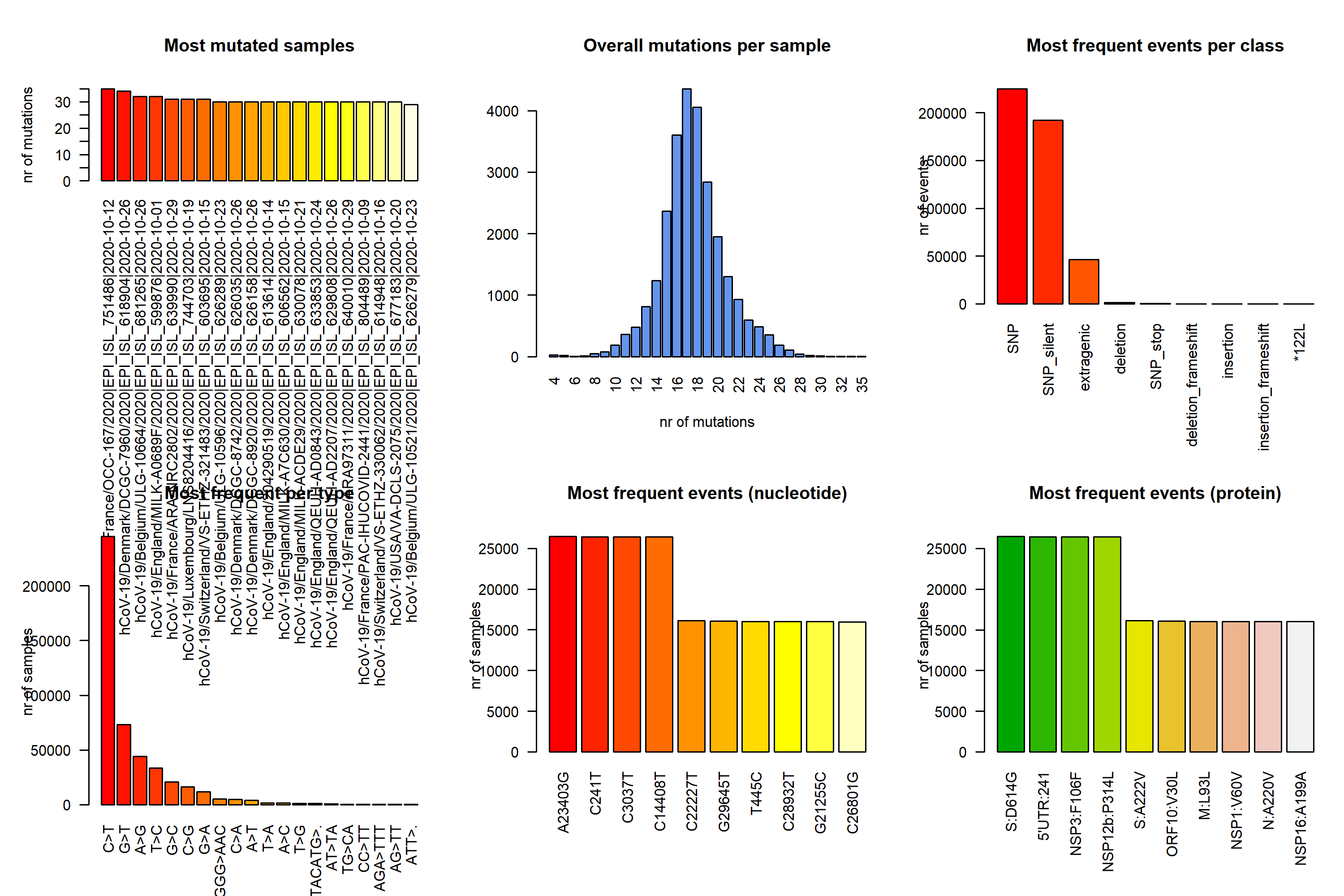

**Figure S11:** Mutational analysis of SARS-CoV-2 genome based on the sequencing data October 2020.

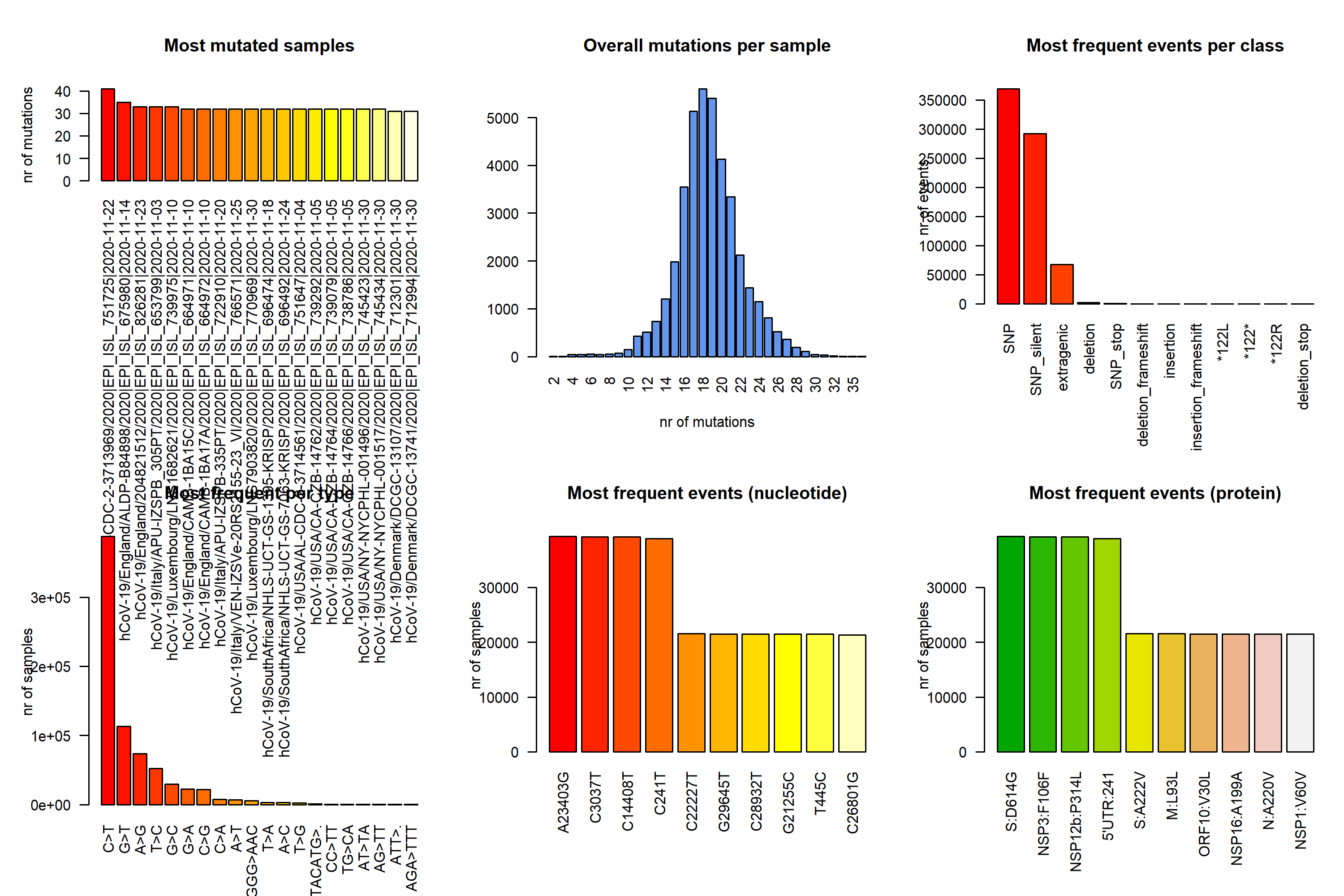

**Figure S12:** Mutational analysis of SARS-CoV-2 genome based on the sequencing data November 2020.

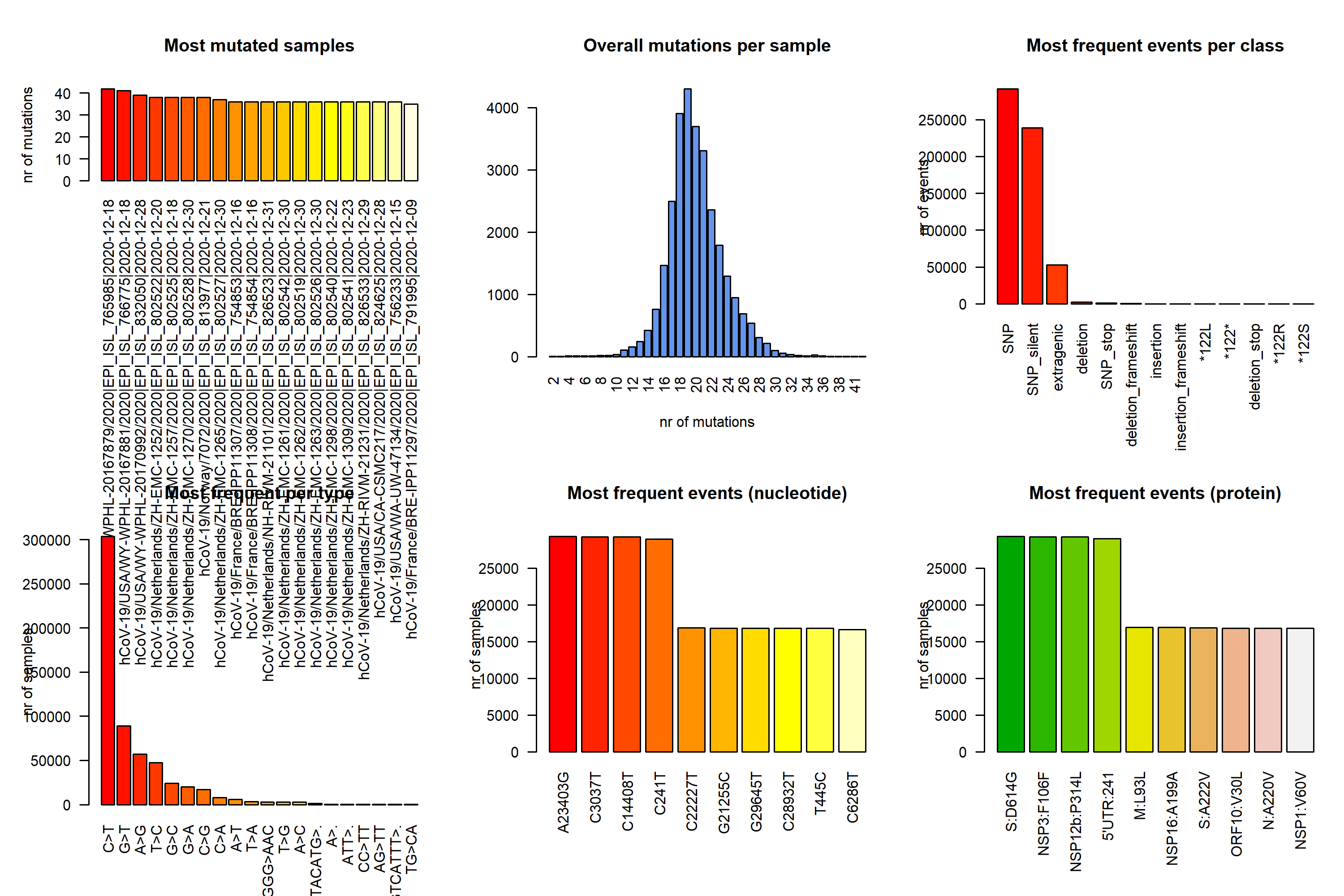

**Figure S13:** Mutational analysis of SARS-CoV-2 genome based on the sequencing data December 2020.
